# The biogenesis and function of nucleosome arrays

**DOI:** 10.1101/2021.02.10.429500

**Authors:** Ashish Kumar Singh, Tamás Schauer, Lena Pfaller, Tobias Straub, Felix Mueller-Planitz

**Affiliations:** Molecular Biology, Biomedical Center, Faculty of Medicine, Ludwig-Maximilians-Universität München, 82152 Planegg-Martinsried, Germany; Bioinformatics Unit, Biomedical Center, Faculty of Medicine, Ludwig-Maximilians-Universität München, 82152 Planegg-Martinsried, Germany; Novartis Institutes for BioMedical Research, CH-4056 Basel, Switzerland; Institute of Physiological Chemistry, Faculty of Medicine Carl Gustav Carus, Technische Universität Dresden, Fetscherstraße 74, 01307 Dresden, Germany

## Abstract

Numerous chromatin remodeling enzymes position nucleosomes in eukaryotic cells. Aside from these factors, transcription, DNA sequence, and statistical positioning of nucleosomes also shapes the nucleosome landscape. Precise contributions of these processes remain unclear due to their functional redundancy *in vivo*. By incisive genome engineering, we radically decreased their redundancy in *Saccharomyces cerevisiae*. The transcriptional machinery is strongly disruptive of evenly spaced nucleosomes, and proper nucleosome density and DNA sequence critical for their biogenesis. The INO80 remodeling complex spaces nucleosomes *in vivo* and positions the first nucleosome over genes in an H2A.Z-independent fashion. INO80 requires its Arp8 subunit but unexpectedly not the Nhp10 module for spacing. Spaced nucleosomes prevent cryptic transcription and protect cells against genotoxic stress such as DNA damage, recombination and transpositions. We derive a unifying model of the biogenesis of the nucleosome landscape and suggest that it evolved not only to regulate but also to protect the genome.

---

Nucleosomes are an ancient innovation of evolution and shape the structure and function of genomes of virtually all eukaryotes. DNA is densely coated with these particles, which profoundly influences access to the underlying genetic information and affects all nuclear processes ^1^.

Nucleosomes arrange on DNA like beads on a string, forming arrays of evenly spaced nucleosomes with a characteristic center-to-center distance, the so-called nucleosome repeat length (NRL). Nucleosome arrays tend to be aligned (“phased”), with respect to the transcription start site (TSS). They are punctuated by nucleosome free regions (NFRs) around promoters. NFRs are important for promoter activity ^2^, and the position of the first nucleosome downstream of the TSS, the +1 nucleosome, helps select the precise start site of transcription^3,4^.

The nucleosome landscape is under constant assault from disruptive processes such as DNA replication and repair ^5^. ATP-dependent nucleosome remodelers are important factors that help to reestablish the nucleosome landscape ^6,7^. The RSC remodeler, for instance, clears the NFR of nucleosomes, and thereby contributes to proper positioning of the +1 nucleosome ^8^. Other remodelers specialize in generating equal spacing between nucleosomes. This activity is quite well documented for ISWI and Chd1 remodelers. Cells lacking these ‘spacing remodelers’ suffer from disrupted nucleosome arrays and closely packed di-nucleosomes ^9–11^.

How spacing remodelers work mechanistically remains contested. Evidence for two scenarios exists. One model posits that spacing remodelers set a characteristic NRL between nucleosomes by ‘clamping’ nucleosomes at fixed distances ^12,13^. The second model proposes that the remodeler measures the length of DNA that flanks the nucleosome, the so-called linker DNA. Long linker DNA activates the remodeler, which then slides the nucleosome efficiently in the direction of the long linker. In doing so, it equilibrates the linker lengths in an array of nucleosomes ^14^.

Nucleosome spacing is difficult to assay *in vitro* due to a dearth of sensitive spacing assays. The movement of a mononucleosome to the center of a short DNA molecule often serves as a proxy of the spacing reaction. ISWI, Chd1 and INO80 are all able to center mononucleosomes on DNA ^15,16^. Consistent with the linker length equilibration model, all these remodelers sensitively react to the length of DNA flanking the nucleosome. INO80, for example, slides nucleosomes ∼100-fold faster when the flanking DNA length increases from 40 bp to 60 bp. Responsible for this switch-like response is its Nhp10 module, which binds to flanking DNA ^17^. The Arp8 module also binds linker DNA and, like Nhp10, contributes to linker DNA sensing^18,19^. It is possible that the ability to sense linker lengths underlies INO80’s ability to position trinucleosomes with ∼30 bp distances^16^.

In principle, evenly-spaced nucleosome arrays could emerge even without the help of remodelers through statistical positioning ^20^. This possibility requires a nucleosome-repellent barrier, which DNA-binding proteins, such as General Regulatory Factors (GRFs), or nucleosome repellent DNA sequences are known to form ^2^. When the nucleosome density is high, nucleosomes downstream of the barrier can only assume a limited number of configurations. From this limited number of configurations, evenly spaced nucleosome arrays emerge in the population average.

A critical test of statistical positioning is reducing the number of nucleosomes. Average distances between nucleosomes should increase, and nucleosome positioning should become fuzzier in the population average. Results from nucleosome depletion experiments in yeast, however, were not fully conclusive. Whereas nucleosome depletion strongly increased fuzziness of nucleosomes, spacing between nucleosomes did not seem to increase sufficiently ^21–23^. Perhaps nucleosomes remain ‘clamped’ together by remodeling enzymes ^12,13^. Studying the function of remodelers in nucleosome-depleted cells therefore promises insights into molecular mechanisms of nucleosome spacing.

In addition to remodelers and statistical positioning, the transcription machinery has been suggested to play pivotal roles in the biogenesis of the nucleosome architecture. The RNA polymerase II preinitiation complex, in conjunction with other factors, could help generate the NFR and position the +1 nucleosome. The +1 nucleosome would then serve as a barrier against which other nucleosomes are packed. In addition, transcription elongation may directly or indirectly establish nucleosome arrays, e.g. through recruiting spacing remodelers ^24,25^. *In vitro* experiments also hint at a role of transcription in organizing the nucleosome landscape. Transcriptionally inactive cell extracts qualitatively establish features of *in vivo* like nucleosome patterns on *in vitro* reconstituted nucleosomes, but not to the degree seen *in vivo* ^26^.

The massive number of processes acting on the nucleosome landscape has hampered previous efforts to disentangle their contributions and study their mechanisms, limiting our understanding of the biogenesis of the nucleosome landscape. Here, we exploit a yeast strain lacking all remodelers of the ISWI and Chd1 family (*isw1Δ, isw2Δ, chd1Δ*; referred to as TKO hereafter) ^27^ to strongly reduce functional redundancy. This strategy allowed us to dissect principles of nucleosome organization *in vivo*. It helped us to reconcile contradictory interpretations with regards to contributions of the DNA sequence, statistical positioning and transcription, and identify INO80 as a nucleosome spacing factor *in vivo*. Of note, we find that INO80 spaces nucleosomes genome-wide, and positions the +1 nucleosome. INO80 does so by relying on the Arp8 but not the Nhp10 subunit or the histone variant H2A.Z. Finally, we provide multiple lines of evidence that nature has evolved spacing remodelers to protect the genome from genotoxic insults. Overall, our results suggest that transcription and nucleosome remodelers compete to establish nucleosome arrays, which helps maintain genome integrity.

## Results

### Nucleosome density and remodelers cooperate to generate nucleosome arrays

Nucleosome arrays are thought to persist after artificial reduction of histones *in vivo*. They appear to possess NRLs that are similar to those of WT cells ^21–23^. We hypothesized that these residual arrays are the product of spacing remodelers of the ISWI and CHD1 family, which we found before to ‘clamp’ nucleosomes at fixed distances *in vitro* ^12^. To test the clamping hypothesis *in vivo*, we employed a histone depletion (HD) system ^28^ in cells lacking *ISW1, ISW2* and *CHD1* (TKO HD). Histone levels were approximately halved compared to WT cells **(Supplementary Fig. 1a, b)**.

Composite plots reveal that the +1 nucleosomes remain well positioned in the TKO HD strain as suggested by a largely comparable peak height of the +1 in WT and TKO HD samples **(Fig. 1a and Supplementary Fig. 1c)**. The +2 nucleosomes are still discernible in TKO HD. Beyond the +2 nucleosomes, however, phased and evenly spaced nucleosome arrays (hereafter simply referred to as regular nucleosome arrays) are largely absent.

**Fig. 1.**
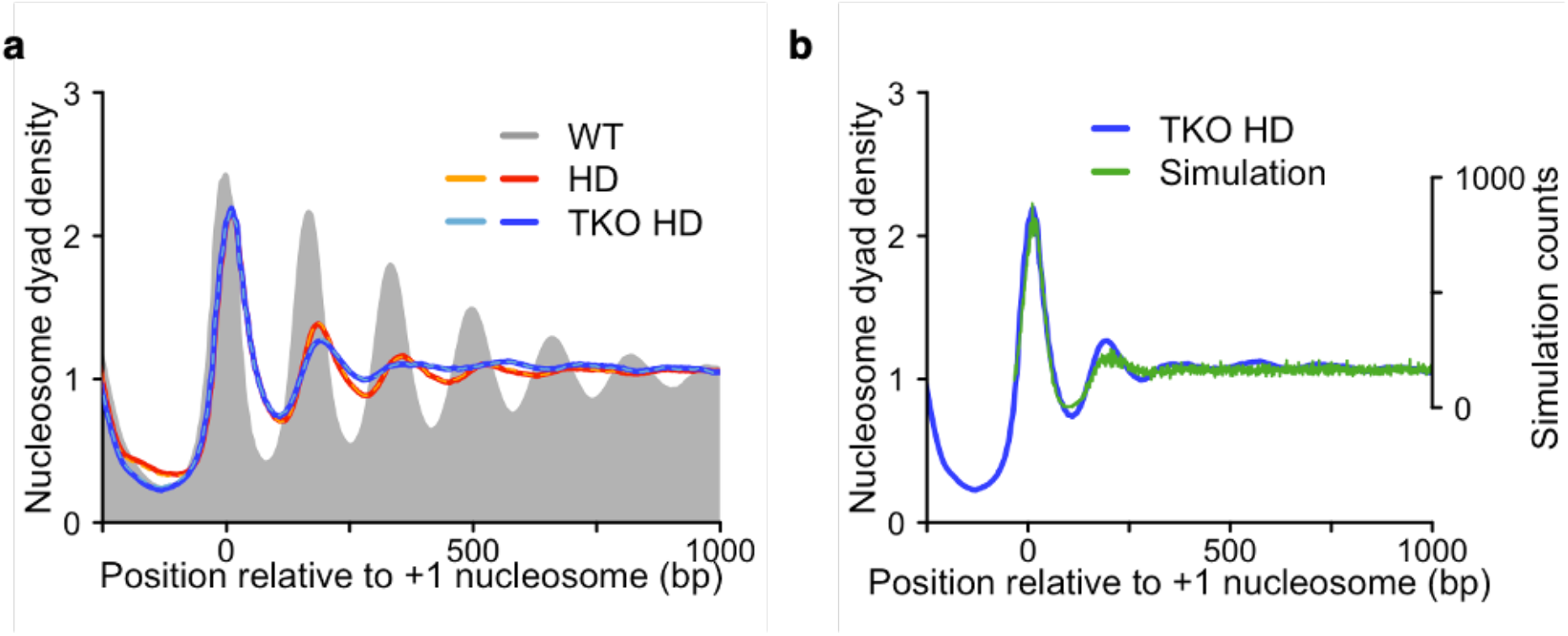
ISWI and Chd1 remodelers rely on proper nucleosome densities to generate nucleosome arrays. **a** Composite plots depicting the average nucleosome organization of ∼5000 genes before (WT; grey) and after histone depletion in otherwise wild-type (HD) or TKO cells (TKO HD). All strains express histones H3 and H4 from a galactose promoter. HD was induced by shifting cells from galactose- to glucose-containing media for 3 h, which reduced histone protein amounts by ∼50% **(Supplementary Fig. 1a, b)**. Nucleosome dyad positions were aligned to known +1 nucleosome positions of WT cells. Dashed lines in brighter color are biological replicates. **b** Simulated, truly random nucleosome organization downstream of a well-positioned +1 nucleosome (green). Nucleosome occupancy was fixed at 51% to simulate histone depletion conditions (see Materials and Methods). TKO HD data from (**a**) is replotted for reference.

We wondered if the +2 nucleosomes in the TKO HD cells are actively positioned by unknown mechanisms or if their positioning emerges from random, i.e. statistical positioning. To test the latter possibility, we simulated random nucleosome positions downstream of a well-positioned +1 nucleosome at a nucleosome density that mimics *in vivo* histone depletion (see Methods). The simulated curve showed a +2 nucleosome at a similar location to the TKO HD sample **(Fig. 1b)**, suggesting that the +2 nucleosomes in the TKO HD sample emerge at least to some extent from statistical positioning.

To test whether spacing remodelers of the ISWI and Chd1 family can induce order in the statistically positioned nucleosomes present in TKO HD cells, we performed HD in otherwise WT cells. Compared to TKO HD, nucleosome arrays in HD cells experienced a modest but consistent increase in amplitude **(Fig. 1a)**. This increase is not due to changes in cell cycle progression as both HD and TKO HD cells enrich to similar extent in the G2/M phase **(Supplementary Fig. 1d)** ^29^.

The results imply that ISWI and/or Chd1 remodelers can order nucleosomes after histone depletion, consistent with a clamping activity ^12^. However, nucleosome arrays seen in HD cells only modestly exceeded that of TKO HD and fell drastically short of in WT cells, **(Fig. 1a)**. We therefore propose that ISWI and Chd1 remodelers cannot overcome the disorder induced by reducing the nucleosome density. High histone density is thus required by ISWI and Chd1 to generate nucleosome arrays.

### Transcription destroys the nucleosome landscape

We noticed that the TKO cells with normal histone levels possess detectable arrays even though the cells are devoid of bona fide spacing remodelers **(Fig. 2a)**^9,10^. Additional factors, e.g. the transcriptional machinery, could be responsible for this residual nucleosome organization ^7,24,25^. If true, one would expect a decline in nucleosome array organization upon transcription inhibition. To test this possibility, we depleted Pol II for 1 h and 2 h in the TKO strain (TKO - Pol II) using anchor-away technology (**Supplementary Fig. 2a)** ^30^.

**Fig. 2.**
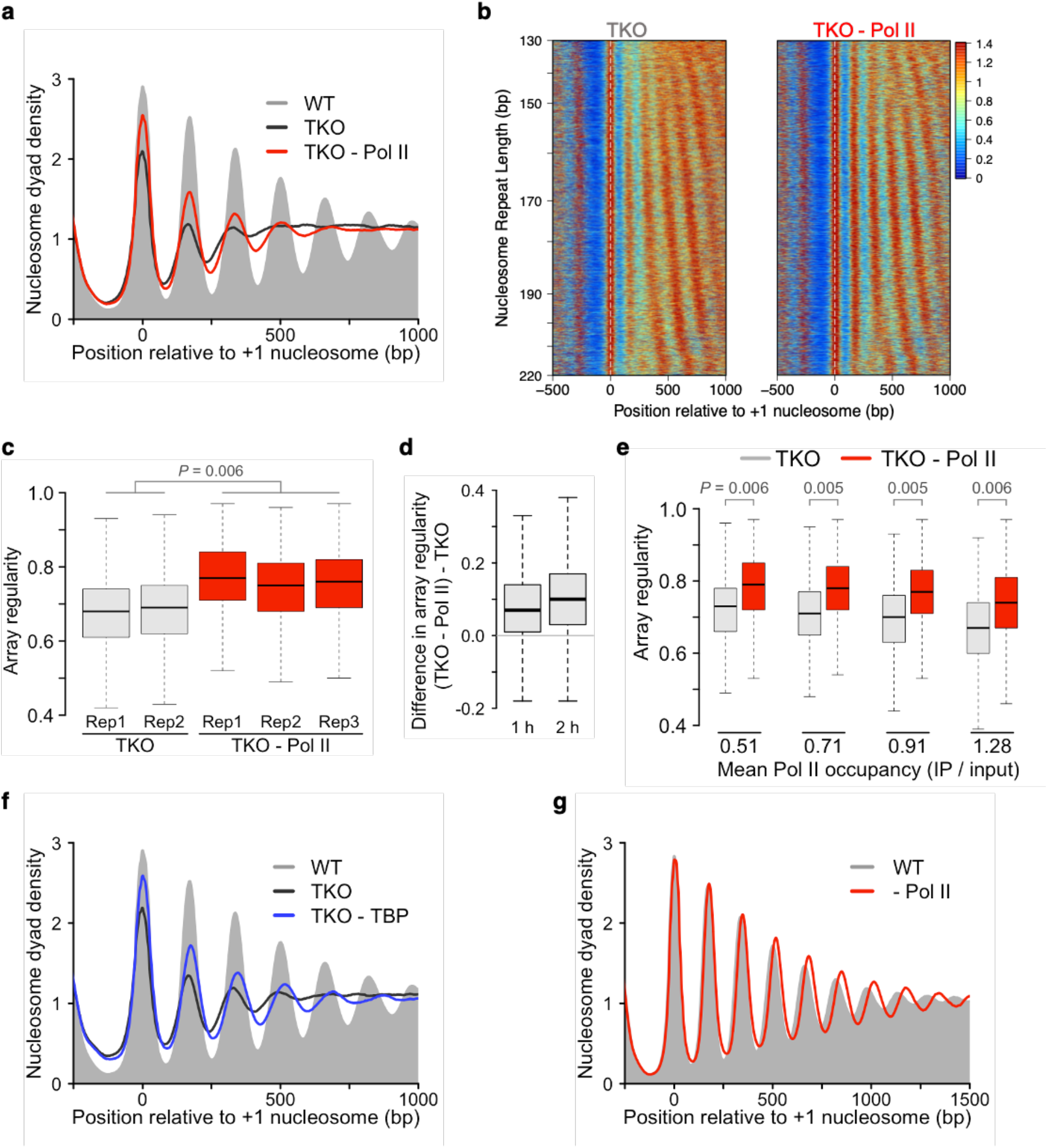
Transcription disrupts the regularity of nucleosome arrays. **a** Pol II depletion in TKO cells using anchor-away technology elevates levels of phased regularly spaced arrays (red line) compared to a TKO control strain (black). A WT control strain is shown for reference (grey). Rpb1 is FRB-tagged in all strains. Cells were either rapamycin- or mock-treated with vehicle for 1 h. **b** Heatmaps for data from (**a**). Genes were sorted according to the NRL observed in TKO cells. Data from two to three biological replicates are pooled. Color scale represents nucleosome dyad density. **c** The median nucleosome array regularity increases in biological replicates (Rep) upon Pol II depletion in TKO cells. Array regularity was determined for each gene by calculating the cross-correlation score between the MNase signal and an idealized Gaussian pattern **(Supplementary Fig. 2e)**. P-value (*P*) represent statistical analyses performed with two-tailed Welch’s t-test on the mean values of individual replicates. **d** Most genes acquire a higher degree of array regularity after Pol II depletion in TKO cells. Shown is the difference in array regularity for each gene before and after Pol II depletion for 1 h and 2 h. **e** Array regularity is rescued upon Pol II depletion irrespective of the transcriptional strength of genes. Genes were sorted by Pol II occupancy and divided into quartiles. Mean Pol II occupancy values of each quartile can be found underneath the plot. Pol II occupancy data for an *isw1Δ, chd1Δ* double mutant strain served as a proxy for TKO cells ^9^. Statistical analysis represents linear mixed-effect model fitted on mean array regularity values of two replicates. **f** Same as (**a**), but for TBP depletion in TKO cells for 1 h. **g** Nucleosome organization upon Pol II depletion in the WT cells for 1 h.

Contrary to the expectation, nucleosome arrays became substantially more pronounced when we blocked transcription **(Fig. 2a, Supplementary Fig. 2b – d)**. To test whether only a subset of genes experiences a rescue of nucleosome arrays upon Pol II depletion, we measured the NRL ^9^ and array regularity ^31^ for each gene **(Supplementary Fig. 2e)**. Heatmaps of the NRL-sorted MNase-seq data reveal that Pol II depletion globally affects the nucleosome landscape and NRL distribution **(Fig. 2b, Supplementary Fig. 2f)**. Average array regularity reproducibly increased over most genes (77.5 % for 1 h and 81 % for 2 h) after Pol II depletion **(Fig. 2c, d)**. The increase in array regularity upon Pol II depletion was genome-wide and not restricted to highly transcribed genes **(Fig. 2e)**, consistent with the pervasive transcription of the genome ^32^.

Pol II depletion shifts cells towards the G1 phase **(Supplementary Fig. 2g)** and could therefore increase array regularity indirectly. We ruled out this possibility by depleting Pol II in fully arrested TKO cells **(Supplementary Fig. 2h, i)**. Other indirect effects remain possible. However, given that we observe a gain of regularity, not a destruction, and that we have identified the factor that causes the increase (see below), we propose that indirect effects are less likely.

The results above leave open the possibility that components of the transcription pre-initiation complex (PIC) can still assemble in the genome and serve as a barrier for phasing arrays ^33^. To test this possibility, we depleted the TATA-binding protein (TBP) in TKO cells. TBP depletion rescued nucleosome arrays **(Fig. 2f)**, similar to Pol II depletion, suggesting that the PIC does not contribute towards establishing regular nucleosome arrays.

The results so far do not support models in which the transcription machinery promotes nucleosome array organization. A caveat remains, however, as transcription may do so only in presence of all remodelers, e.g. by recruiting them to chromatin. In contrast to the prediction of this scenario, the nucleosome array regularity was not compromised upon depletion of Pol II in WT cells **(Fig. 2g, Supplementary Fig 2a)**. In fact, in six available datasets collected in three laboratories, array regularity increased, not decreased, after Pol II depletion **(Supplementary Fig 3b – e)** ^34,35^. We therefore suggest that the act of transcription is disruptive to nucleosome arrays even in presence of spacing remodelers.

### The INO80 remodeling complex induces spacing of nucleosomes *in vivo*

The rescue of array regularity upon Pol II depletion in TKO cells was intriguing because it implies the existence of another spacing factor in addition to the bona fide spacers of the ISWI and Chd1 family. The INO80 complex is an attractive candidate because its deletion in WT cells measurably changes array structure *in vivo* ^23,36^, and it can generate short arrays with three nucleosomes *in vitro* ^16,26^.

To test whether the INO80 complex is responsible for the rescue upon Pol II depletion, we co-depleted INO80 and Pol II in TKO cells (**Supplementary Fig. 2a, 4a**). Unlike Pol II single depletion, co-depletion of Pol II and INO80 did not increase the regularity of nucleosome arrays **(Fig. 3a and Supplementary Fig. 4b, c)**, suggesting that INO80 is the responsible factor that forms arrays after Pol II depletion. Consistent with an array forming activity of INO80, its depletion in actively transcribing TKO cells further decreased nucleosome arrays and array regularity **(Fig. 3b and Supplementary Fig. 4d – e)**. We ruled out that these effects were confined to genes that suffer from an altered Pol II distribution upon INO80 depletion **(Supplementary Fig. 4f, g)** or to genes with a ChEC-seq signal for INO80 (not shown) ^3^, suggesting that INO80 works throughout most of the genome. The combined results imply that INO80 helps generate regular nucleosome arrays genome-wide in TKO cells, and that it can do so in presence or absence of active transcription.

**Fig. 3.**
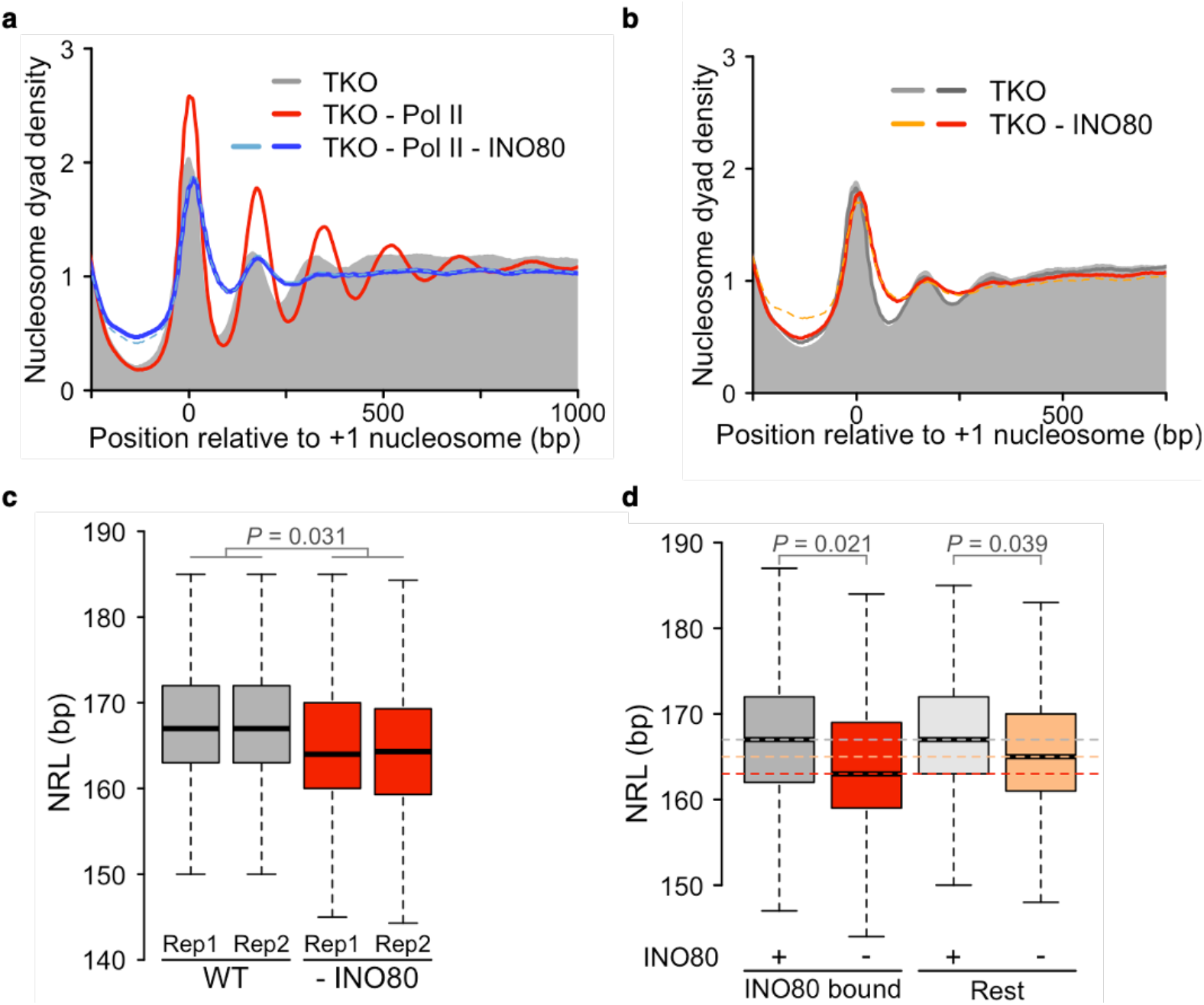
INO80 induces nucleosome array formation *in vivo*. **a** Gene-averaged nucleosome organization for the indicated Pol II anchor-away strains. Samples were treated with vehicle (TKO) or rapamycin (all other samples) for 2 h, twice as long as in Fig. 2. The treatment regimen allowed for efficient depletion, as tested by live cell imaging of GFP-tagged INO80 and Rbp3 ChIP-qPCR **(Supplementary Fig. 2a, 4a)**. Dashed line: biological replicate. **b** Nucleosome organization upon INO80 depletion (1.5 h) in TKO cells (red). Ino80-GFP-FRB tagged TKO cells treated with vehicle served as the reference (grey). Live cell imaging confirmed INO80 depletion **(Supplementary Fig. 4d). c** NRL distribution before and after INO80 depletion (1.5 h) in WT cells. Median values are at 167 bp and 164 bp for WT and INO80 depleted cells, respectively. **d**, NRLs of 1646 genes bound by INO80 (median value of 163 bp) as measured by ChEC-seq ^3^ compared to rest of the genes (165 bp).

To test whether INO80 contributes to the nucleosome landscape also in WT cells, we depleted INO80. Depletion of INO80 diminished nucleosome regularity, albeit to varying degrees in our and published dataset **(Supplementary Fig. 4h, i)**. Importantly, depletion consistently reduced the genome-wide NRL by 3 bp in our and published data ^37^ **(Fig. 3c, Supplementary Fig. 4j)**. INO80-bound genes ^3^ experienced a larger decrease (4 bp) compared to all other genes (2 bp) **(Fig. 3d)**. An alteration in cell cycle progression upon INO80 depletion ^38^ is unlikely to contract the NRL because the NRL distribution does not change during cell cycle **(Supplementary Fig. 4k)** ^39^. Overall, these results suggest that INO80 contributes to NRL determination in WT cells.

### The Arp8, not the Nhp10 module regulates INO80 spacing in vivo

Our ability to isolate INO80 activity provided us with the opportunity to dissect the spacing mechanism of INO80 *in vivo* **(Fig. 4a)**. The Nhp10 module of INO80 imparts a switch-like response on the sliding activity in response to small changes in the linker length ^17^. As such, the Nhp10 module is a strong candidate to control nucleosome spacing. To test this possibility *in vivo*, we deleted *NHP10* from the TKO - Pol II system. While the amplitude of arrays modestly reduced upon *NHP10* deletion **(Supplementary Fig. 5a)**, the NRL distribution with and without Nhp10 remained similar **(Fig. 4b)**. Comparable effects were observed after *NHP10* deletion in transcribing WT and TKO cells **(Supplementary Fig. 5e, g, k)** ^40^ and after deletion of the 300 N-terminal amino acid residues of the Ino80 ATPase subunit **(Supplementary Fig. 5f, h, k**). These residues contribute to the association of the Nhp10 module to INO80 and proteolytically degrade when Nhp10 is missing ^17,41^ **(Supplementary Fig. 5i**). Concurrent deletion of *NHP10* and the N-terminus of Ino80 in WT and TKO cells also showed negligible effects (**Supplementary Fig. 5k** and data not shown). Collectively, the results challenge the hypothesis that the Nhp10 module or the N-terminus of Ino80 are critically required for the INO80 complex to evenly space nucleosomes and to determine INO80’s preferred genome-wide NRL.

Like Nhp10, Arp8 has been implicated in linker DNA sensing ^18,19^. When we deleted *ARP8*, INO80 lost most of its ability to evenly space nucleosomes in Pol II-depleted TKO cells **(Supplementary Fig. 5b)**, attesting to Arp8’s important catalytic role ^42^. Nevertheless, the residual arrays generated in absence of Arp8 had an NRL distribution that was shifted to shorter values, peaking at 151 bp instead of the 167 bp observed in presence of Arp8 **(Fig. 4c)**.

The shift to smaller NRL upon *ARP8* deletion can be observed also in WT and TKO cells **(Fig. 4d, e and Supplementary Fig. 5c, j, k)**. The NRL distribution in *arp8Δ* cells could be faithfully rescued by expression of wild-type Arp8. Notably, Arp8 lacking 197 N-terminal amino acids, which interact with the linker DNA^18^, did not rescue the phenotype (**Fig. 4e**). In conclusion, these results highlight the importance of Arp8 and its N-terminal region for INO80-mediated remodeling and suggest that Arp8 helps INO80 to determine the NRL.

**Fig. 4.**
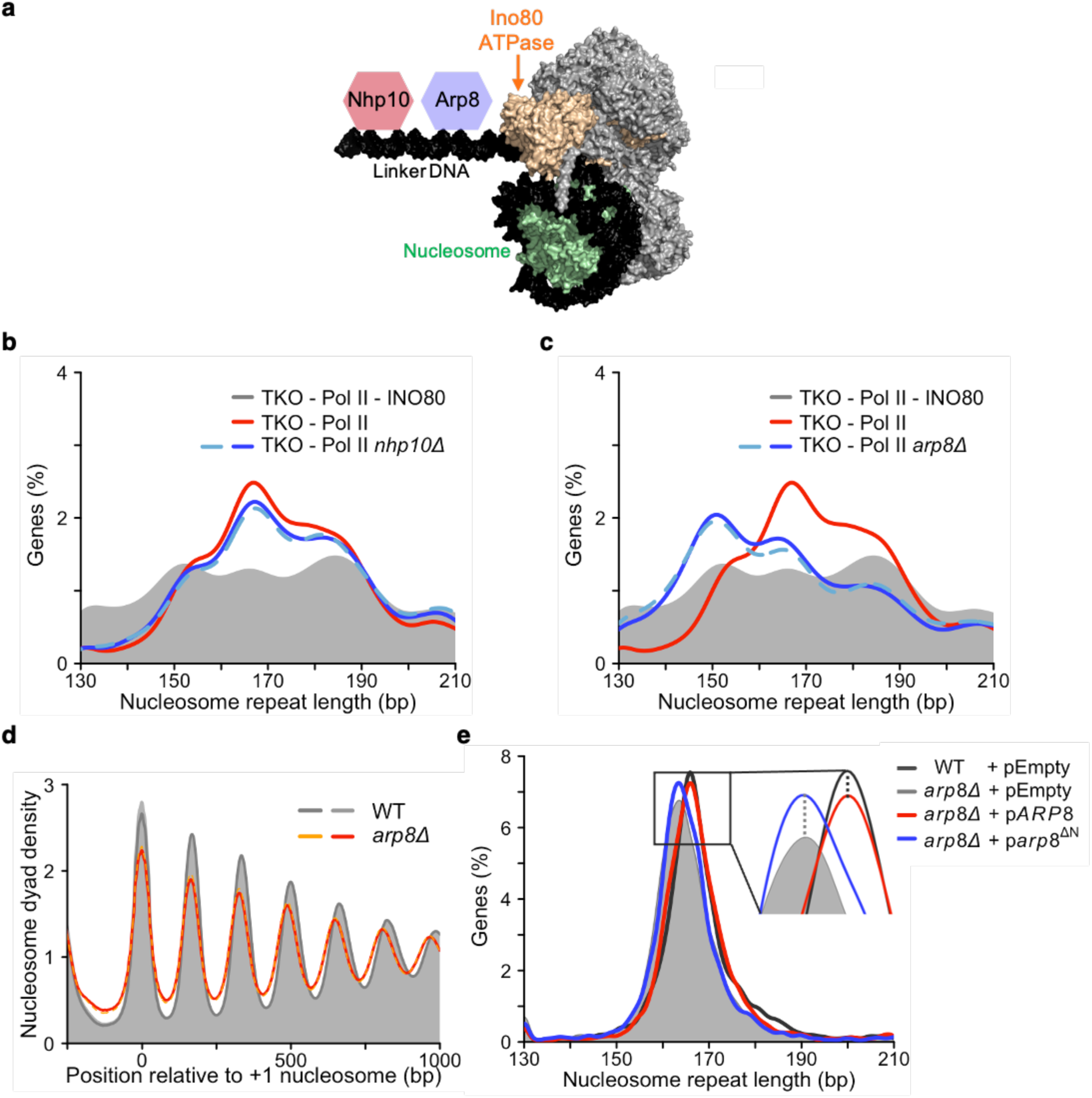
Arp8, but not Nhp10, regulates INO80-mediated spacing *in vivo*. **a** Cartoon of the INO80 complex bound to the nucleosome based on PDB code 6FML. The Nhp10 and Arp8 modules interact with linker DNA. **b** Deletion of *NHP10* (blue lines) does not substantially alter the NRL distribution in TKO -Pol II samples (red; all colored lines peak at 167 bp). Cells depleted for INO80 are shown as a reference (grey). **c** Same as b, but for *arp8Δ*. The NRL distribution shifts left with a peak at 151 bp. **d** Nucleosome organization in WT and *arp8Δ* cells. Deletion of *ARP8* decreases array regularity and NRL. **e** Deletion of *ARP8* in otherwise WT cells shifts the NRL distribution to shorter values (peaks at 163 bp for *arp8Δ* and 166 bp for WT cells). This effect can be faithfully rescued by expression of wild-type Arp8 but not Arp8 lacking amino acids 2-197 (dashed lines). pEmpty: empty vector.

### INO80 positions the +1 nucleosome independently of H2A.Z in vivo

The results above indicate that INO80 helps increase the levels of phased regular arrays. It could do so by spacing nucleosomes in gene bodies and/or by positioning the +1 nucleosome. To test the latter, we inspected the signal heights of the +1 nucleosome. We noticed that the +1 peak was more pronounced after Pol II depletion in TKO cells in composite plots, and that this increase depended on the presence of INO80 **(Fig. 3a)**. To substantiate this observation, we measured +1 nucleosome peak intensities at each gene ^43^. Peak intensities of most +1 nucleosome increased after Pol II depletion in an INO80 dependent manner **(Fig. 5a)**. Moreover, +1 peak intensities reduced upon INO80 depletion in TKO cells **(Fig. 5b)**. We therefore suggest that the INO80 complex has a role in positioning the +1 nucleosome *in vivo*, in line with previous observations^26,37^.

**Fig. 5.**
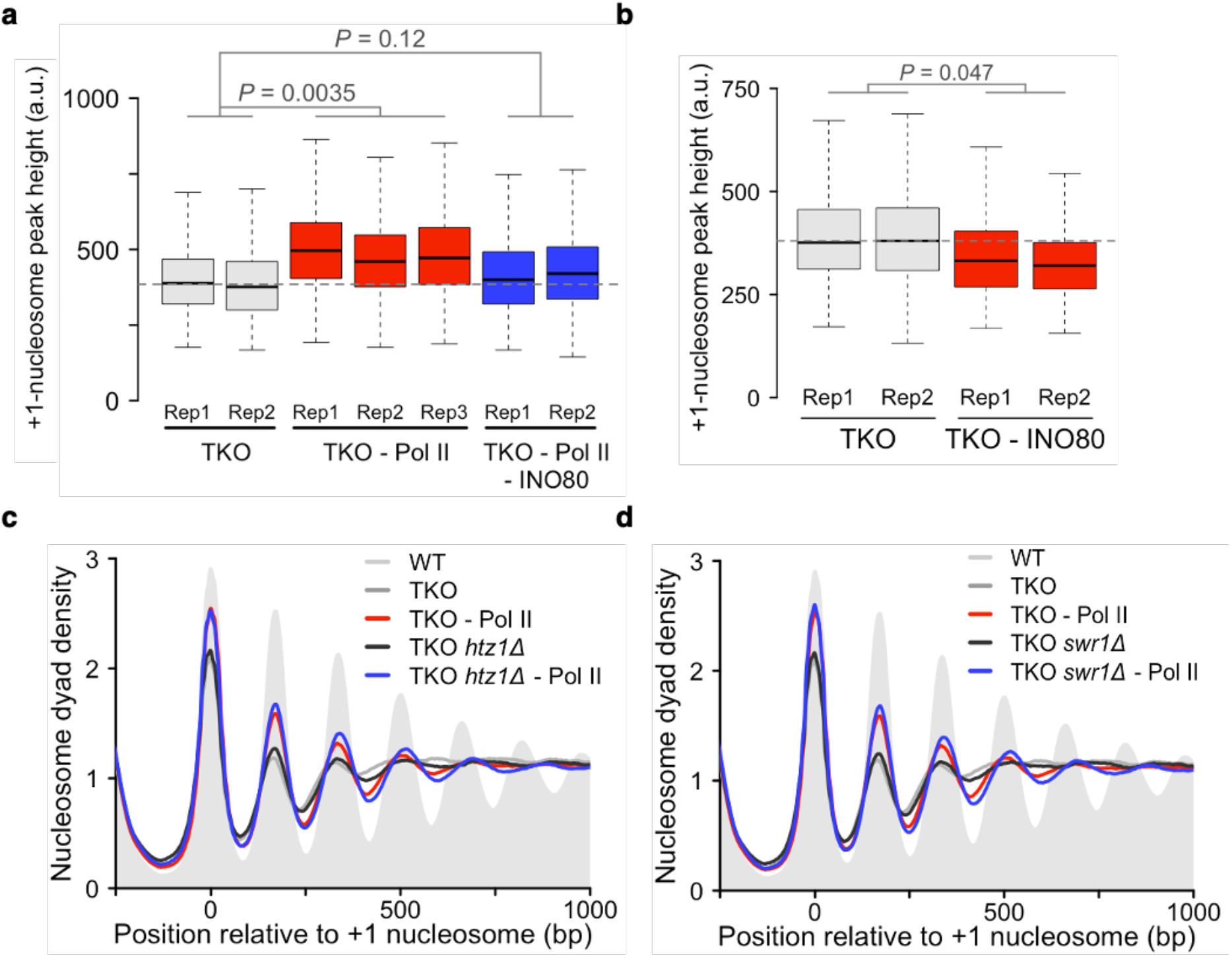
IN080 positions +1 nucleosomes independently of H2A.Z. **a** Box plots of peak heights measured for the +1 nucleosomes of ∼5000 genes in the indicated strains and replicates. TKO is a Rpb1-FRB tagged strain treated with vehicle for 1 h. The horizontal dashed line indicates the mean of two biological replicates of the TKO strain. P-values (*P*) represent statistical analyses performed with one-way Anova followed by post-hoc Tukey HSD test on the mean values of individual replicates. **b** Same as (**a**), but for INO80 depletion in TKO. The TKO reference is Ino80-GFP-FRB tagged cells treated with vehicle. Statistical analysis was performed with two-tailed paired Welch’s t-test on the mean values of two replicates. **c** Deletion of H2A.Z (*htz1Δ*; blue) has no deleterious effects on +1 peak heights and nucleosome arrays upon Pol II depletion (red). The negative controls (grey and black) are the corresponding vehicle-treated Rpb1-FRB tagged strain. Pol II was depleted for 1 h. **d** Same as (**c**), but for *swr1Δ*. Data in (**c**) and (**d**) are averages of two or three biological replicates.

The +1 nucleosome is strongly enriched with the histone variant H2A.Z, particularly upon Pol II depletion ^34^. INO80 slides H2A.Z-containing nucleosomes two- to four-fold faster than canonical nucleosomes *in vitro* ^44,45^, leading to a simple model that INO80 recognizes H2A.Z and thereby positions the +1 nucleosomes particularly well. Strikingly, H2A.Z deletion had no adverse effect on the peak height and width of the +1 nucleosome in TKO cells, even after Pol II depletion where INO80 activity can be visualized best **(Fig. 5c)**. We independently validated these results by deleting the SWR1 remodeler, which deposits H2A.Z at +1 nucleosomes **(Fig. 5d)**. We conclude that INO80 positions +1 nucleosomes independently of H2A.Z.

### The DNA sequence affects the NRL

The location of the +1 nucleosome is encoded in part by nucleosome-attracting DNA sequences ^46^. Close inspection reveals that +1 nucleosomes tend to reside 17 bp downstream of the thermodynamically most preferred positions **(Table 1)**. Depletion of RSC ^47^ returns the +1 nucleosomes to within 1 bp of the nucleosome-attracting DNA sequences. When cells lacked INO80, the +1 nucleosomes moved into the gene body, further away from the thermodynamically favored position **(Table 1)**. This effect has been observed before and agrees with the suggestion that INO80 and RSC engage in a tug-of-war over the positions of the +1 nucleosomes ^3^.

**Table 1.**
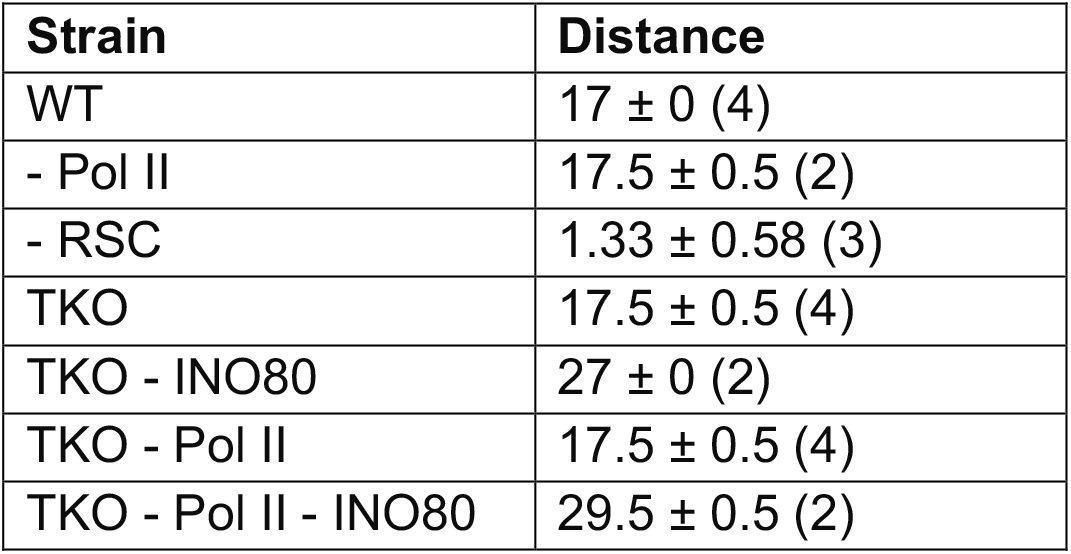
Genome-averaged distances between the +1 nucleosome peaks in MNase-seq data and +1 nucleosome positions predicted from nucleosome affinities. Values are mean and SD of replicates (values in brackets: number of replicates). MNase-seq data for RSC-depleted cells from ^47^.

We next tested whether DNA sequences influence the NRL. Remodelers are believed to override DNA sequence-encoded nucleosome positions and impose a remodeler-encoded NRL ^25^. If true, the observed NRL should not be influenced by DNA sequence.

We calculated nucleosome affinities **(Fig. 6)** ^48^ and divided into quartiles after sorting the genes by their MNase-seq derived NRL for WT cells. Peaks emerged after averaging nucleosome affinities in all quartiles **(Fig. 6a, d)** and faithfully tracked the measured NRL in three quartiles (Q2 to Q4). Similar trends were observed with nucleosome affinities calculated by a different strategy **(Supplementary Fig. 6c)**^46^. The results suggested that DNA sequence contributes to the NRL in much of the genome.

**Fig. 6.**
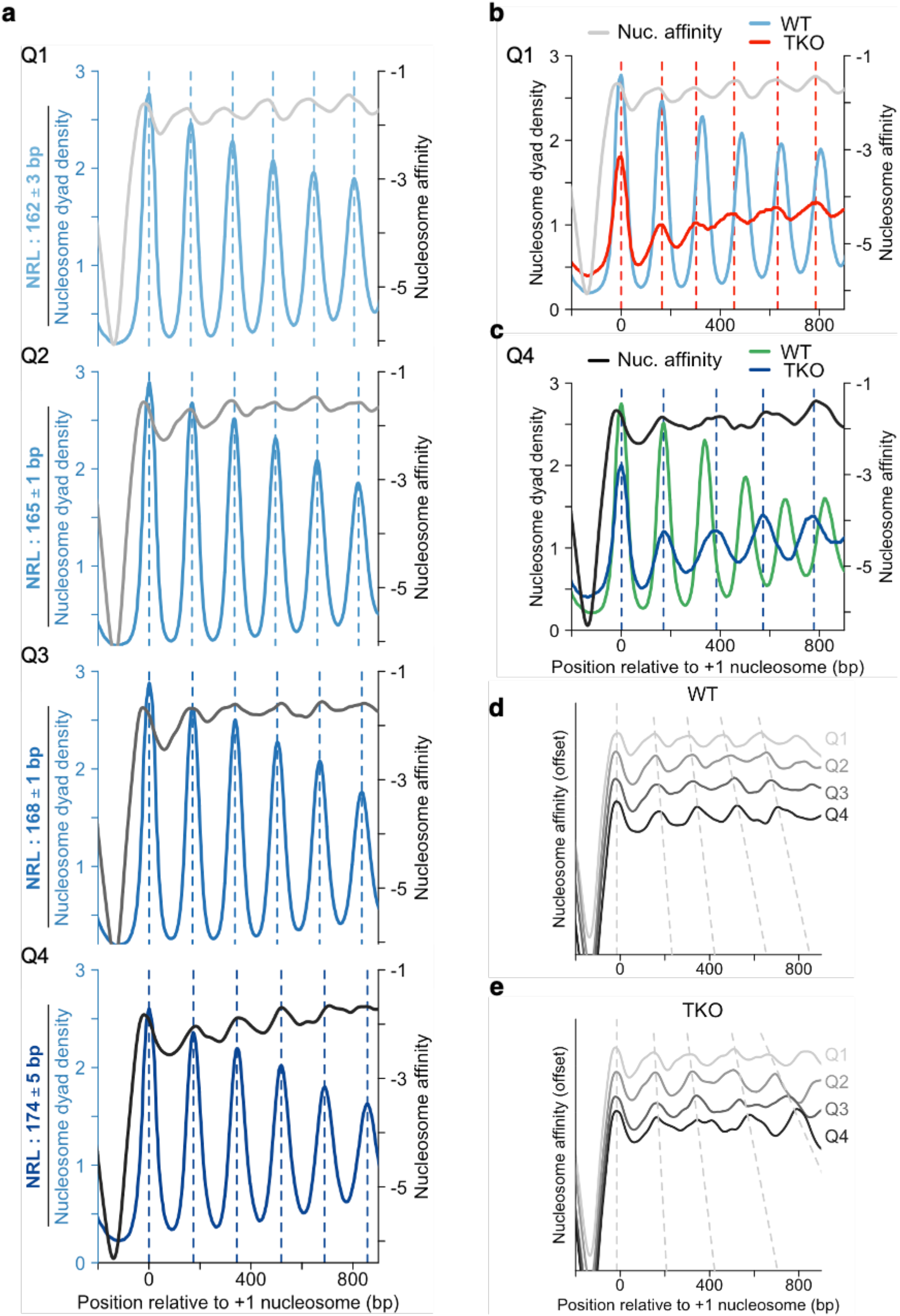
DNA sequence contributes to NRL determination. **a** Comparison of gene-averaged MNase-seq data and calculated nucleosome affinities for WT cells. Genes were filtered (see Methods), sorted by NRL and grouped into quartiles from Q1 (shortest) to Q4 (longest NRL). NRL values are mean NRLs and their SD. Nucleosome affinities were calculated using nuCpos ^48^. **b** Arrays in Q1 of WT cells assume a shorter NRL in TKO cells. This shorter NRL is more similar to the NRL predicted from nucleosome affinities. **c** The NRL of arrays in Q4 of TKO cells is broadly consistent with the NRL predicted from nucleosome affinities. The same genes assume a substantially shorter NRL in WT cells, with peaks in the WT MNase-seq data straying further away from the thermodynamically most stable positions. **d, e** Nucleosome affinities for Q1 – Q4 of WT (**d;** replotted from **a**) or TKO cells (**e**).

Only in Q1 with smallest NRL peaks in nucleosome affinity and MNase-seq did not overlap. The thermodynamically preferred NRL in Q1 was even shorter than the measured one. Of note, the predicted and measured NRLs of the same genes in Q1 matched much better in TKO than in WT cells **(Fig. 6b)**. We therefore propose that ISWI and/or Chd1 remodelers override short, DNA-encoded NRLs and increase their NRL.

TKO cells also possess unusually long NRLs ^9^. To test whether they are at least in part DNA-sequence encoded, we superimposed MNase-seq data of Q4 from TKO cells with calculated nucleosome affinities. Peaks overlapped well. In contrast, WT MNase-seq data for the same set of genes showed smaller NRL **(Fig. 6c)**. We therefore suggest that ISWI and/or Chd1 remodelers drastically shorten long, DNA-sequence encoded NRLs. By decreasing long and increasing short NRLs, these remodelers conceivably contribute to equalizing linker lengths across the genome.

From these results one may expect that arrays in remodeler-depleted cells are more strongly DNA-sequence encoded than in WT cells. Consistent with this expectation, peaks of MNase-seq and nucleosome affinity data overlapped reasonably well in all quartiles for TKO and TKO - Pol II cells **(Fig. 6e, Supplementary Fig. 6a, b, d)**. The amplitudes of calculated nucleosome affinities however do not become larger than WT as might be expected. We therefore propose that the thermodynamic landscape provided by the DNA sequence is relatively shallow with many potential locations for nucleosomes, such that nucleosomes tend to find thermodynamically preferred positions.

### Regular nucleosome arrays affect transcription and protect the genome

With ISWI, Chd1 and INO80 remodelers, the cell has evolved three families of spacing remodelers capable of generating nucleosome arrays. The functional relevance of nucleosome spacing, however, remains unknown. We hypothesized that arrays maintain genomic integrity by protecting the underlying DNA from external insults.

First, we checked whether array regularity anticorrelates with susceptibility to DNA damage. We tested the stress response in cells that lack combinations of remodelers against zeocin, methyl methanesulfonate (MMS), hydroxyurea (HU) or ultraviolet (UV) radiation. Zeocin is a radiomimetic agent, that generates free radicals and induces double-strand (ds) breaks. MMS and HU induce genome instability by inhibiting replication fork progression and depleting dNTPs, respectively ^49^, whereas UV induces formation of pyrimidine dimers. We found that combined loss of ISWI and Chd1 remodelers made cells highly susceptible to zeocin, MMS and HU but not to UV **(Fig. 7a, Supplementary Fig 7a)**. The growth defects of the mutant strains largely anticorrelated with the average array regularity of those cells, consistent with the notion that regular nucleosome arrays protect against DNA damage. Cells lacking *ARP8* were an outlier under HU and MMS stress as Arp8 has additional roles during replication and DNA repair ^38,50^.

**Fig. 7.**
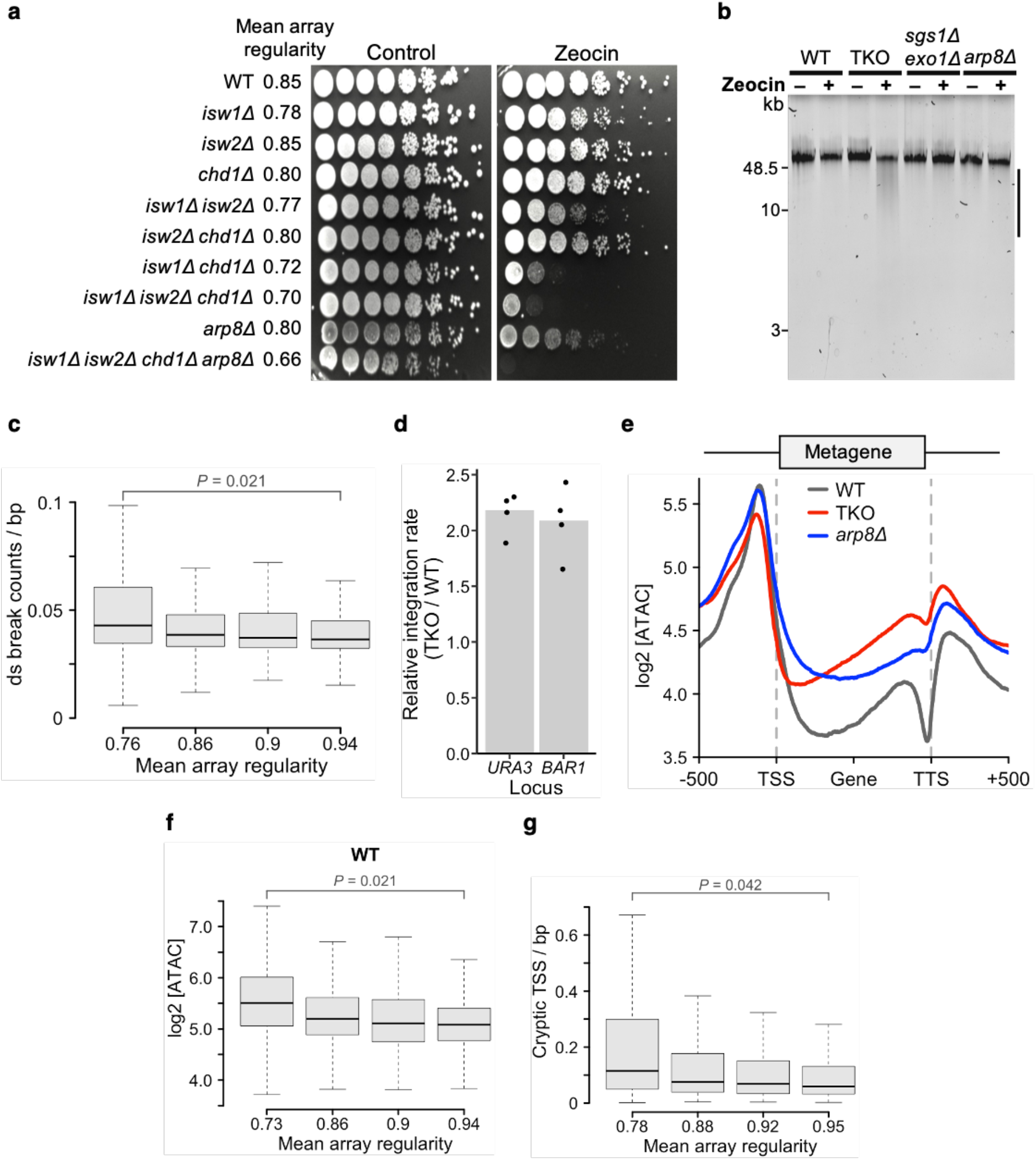
ISWI and Chd1 remodelers protect the genome from genotoxic stress. **a** Growth assay for indicated yeast strains on YPAD with and without Zeocin (100 µg/ml). **b** Zeocin-induced fragmentation of genomic DNA (bar) for the indicated strains. Cells were treated with deionized water (-) or Zeocin (+; 1 mg/ml) for 10 min. **c** Spo11-induced DNA ds breaks ^69^ enrich in genes with low array regularity. **d** The TKO strain endures higher levels of ectopic recombination than the WT. Homologous recombination tested at two genomic loci (*URA3, BAR1*). Dots represent individual replicates from four independent experiments. **e** Metagene plot of ATAC-seq signals for WT, TKO and *arp8Δ* cells. **f** The number of ATAC-seq insertions into gene bodies anti-correlates with array regularity in WT cells. Genes were filtered for an absolute nucleosome occupancy between 0.78 and 0.88 ^61^, sorted by array regularity measured in WT cells and then divided into four quartiles. **g** Cryptic TSSs are enriched in gene bodies with low array regularities. TSS data from ^32^. P-values in (**c, f, g)** represent statistical analyses performed with two-tailed paired Welch’s t-test on the mean values of two replicates.

Strikingly, 10 min of zeocin exposure were enough to visibly degrade genomic DNA in TKO cells but not in WT and *arp8Δ* cells. Cells defective in ds break repair (*sgs1Δ exo1Δ*) also did not show increased levels of breaks, suggesting that increased ds breaks in TKO cells is not derived from defective DNA repair in 10 mins of zeocin exposure **(Fig. 7b and Supplementary Fig. 7b)**. Of note, genes covered with more regular nucleosome arrays are also less likely to suffer from naturally occurring DNA ds breaks during meiosis in WT cells **(Fig. 7c and Supplementary Fig. 7c)**. These observations suggest that spacing remodelers may have evolved in part to mitigate the dangers of DNA damage.

Second, we tested if even spacing between nucleosomes prevents ectopic recombination of DNA into the genome. We monitored homologous recombination rates at two genomic loci ^51^. Transgenes recombined ∼two-fold better in TKO compared to WT cells **(Fig. 7d)**. The TKO mutation also resulted in elevated recombination rates in *arp8Δ* cells **(Supplementary Fig. 7d)**, even though *arp8Δ* show ∼3-fold lower recombination rates than WT ^51^. In summary, ISWI and Chd1 factors tend to suppress recombination. An attractive model is that they do so by keeping the genome evenly coated with nucleosomes, thus limiting DNA accessibility. INO80 on the other hand promotes recombination^51,52^, likely via mechanisms that go beyond formation of arrays.

Third, we followed up on the hypothesis that spacing remodelers limit DNA accessibility. If true, transposon integration rates would be elevated in cells with more irregular nucleosome arrays. To measure transposon integration rates, we performed ATAC-seq in WT, TKO and *arp8Δ* cells. Consistent with the model, gene bodies in TKO cells received substantially higher levels of ATAC integrations than WT cells **(Fig. 7e)**. Cells lacking *ARP8* showed an intermediate phenotype, consistent with intermediate levels of disruption of nucleosome arrays in these cells **(Fig 7e, Supplementary Fig. 7e)**. Intriguingly, ATAC integration rates over gene bodies anti-correlated with measured nucleosome regularity in WT cells (**Fig. 7f**). This effect was detectable, albeit with lower statistical significance, in *arp8Δ* **(Supplementary Fig. 7f)** and TKO cells (not shown). A similar correlation between transposition events and irregularity of nucleosome arrays also holds *in vivo* in WT cells **(Supplementary Fig. 7g)** ^53^.

Lastly, we interrogated whether irregular nucleosome arrays could give rise to cryptic transcription by analyzing published TSSs dataset in WT cells ^32^. We found that genes covered with more regular nucleosome arrays tend to harbor fewer cryptic TSSs (**Fig. 7g**). A genome-wide correlation between average nucleosome regularity and cryptic transcription also holds across mutants that possess disrupted nucleosome arrays, for example TKO cells and cells with defects in histone chaperones such as Spt6 and FACT ^23,54^. We thus suggest that spaced nucleosome arrays contribute to preventing cryptic transcription arising from the gene body.

## Discussion

The nucleosome landscape is shaped by numerous nuclear processes including a variety of nucleosome remodeling complexes and the transcription machinery. It is an experimental challenge to cleanly disentangle the effects of any one of these actors on the nucleosome architecture due to their strong functional redundancy. Here we radically diminished redundancy of factors implicated in the biogenesis of the nucleosome landscape by simultaneously deleting or depleting Chd1, ISW1, ISW2, INO80, and components of the transcription machinery. This strategy helped us to isolate the role of transcription and led to the discovery that the INO80 remodeler helps shape the canonical nucleosome architecture genome-wide.

### The mechanism of INO80-mediated spacing

We capitalized on our ability to isolate INO80 activity in the physiological environment of a cell to study its function and dissect its mechanism, an endeavor that traditionally is carried out more often *in vitro*. We find that the INO80 complex helps position the +1 nucleosome and space nucleosome arrays over genes **(Fig. 3 and Fig. 5a)**. It does so throughout the genome.

How does INO80 accomplish nucleosome spacing? In the simplest model, INO80 positions only the +1 nucleosome. This well-positioned nucleosome then serves as a barrier for randomly positioned downstream nucleosomes. Arrays would then emerge over genes via statistical positioning. In an alternative but mutually not exclusive model, INO80 can actively space arrays. Our Arp8 results suggest that it can in principle. Upon *ARP8* deletion, the NRL sharply decreases **(Fig. 4c)**. This observation is consistent with a role of the Arp8 module in linker-length sensing but cannot readily be explained by statistical positioning theory. Taken together, we suggest that INO80 can perform nucleosome spacing itself, a conclusion that is further substantiated by observations *in vitro* ^16,26^. The results extend the family of bona fide spacing remodelers to INO80, in addition to ISWI and Chd1 remodelers.

Why does INO80 need Arp8 for spacing? Arp8, conserved from yeast to mammals, is thought to sense the presence of linker DNA ^18^. Deprived of this ability, Arp8-less Ino80 may not be able to space nucleosomes any longer ^18^ which could contribute to lower overall nucleosome regularity in *ARP8*-lacking cells **(Fig. 4c, d)**. The yeast-specific Nhp10 module, on the other hand, is not a critical component for INO80-mediated spacing **(Fig. 4b)**. We could imagine, however, that yeasts evolved Nhp10 to optimize the efficiency of INO80-mediated spacing or help INO80 to space arrays in special situations, for example during replication or DNA damage ^55^.

### Transcription is disruptive to nucleosome arrays

Active transcription has been suggested to be critical for the biogenesis of phased regular nucleosome arrays over genes ^25^. Our data instead supports an overall disruptive effect **(Fig. 2)**. The disruptive effect may also be conserved in higher organisms because array regularity tends to be lower in highly transcribed genes ^56^. We propose that spacing remodelers counteract the disruptive effect of transcription. The opposing activities from nucleosome-organizing and -disrupting factors could provide opportunities for chromatin-based regulation.

How transcription destroys array regularity remains unclear. One model posits that elongating Pol II evicts histones ^57,58^. Several considerations argue against this model. First, we observe disruption of nucleosome regularity by the transcription machinery throughout the genome, not only at heavily transcribed genes **(Fig. 2b, e)**. Second, loss of histones appears to precede transcription, not be caused by it^59,60^.

Third, nucleosome occupancy over genes does not correlate with their transcriptional activity ^61^. A model that would be consistent with available data invokes the elongation machinery to reposition nucleosomes. Indeed, we and others observed increase in NRLs upon Pol II depletion **(Supplementary Fig. 3a)** ^34,62^, suggesting that the transcription machinery moves nucleosomes upstream of their original locations. Such an activity, if not counteracted by remodelers, would over time destroy array regularity.

### Biogenesis of the nucleosome landscape

Besides transcription, DNA replication and damage are disruptive to the nucleosome landscape ^5^. Our findings extend previous models how cells reestablish the nucleosome landscape ^24,26^.

We envision four contributing principles that jointly sculpt the nucleosome landscape^63^. First, nucleosomes preferentially accumulate on DNA sequences that inherently possess a high affinity to nucleosomes. We observe such an accumulation in three quarters of the WT genome and in remodeler- and histone-depleted cells. Despite drastically different nucleosome organization in these mutant cells, we did not observe nucleosomes to become substantially enriched or depleted from sites with high nucleosome affinity **(Supplementary Fig. 6a, b)**. We thus suggest that the nucleosome affinity landscape flexibly allows for several thermodynamically equivalent nucleosome configurations.

A second process helps form the NFR. Nucleosome destabilizing DNA sequences enriched at the NFR, GRFs and the RSC remodeler play important roles in keeping nucleosomes away from NFRs ^8^.

A third process positions the +1 nucleosome. RSC pushes the +1 nucleosome away from its highest affinity site further into the gene body **(Table 1)**. The +1 nucleosome continues to slide deeper into the gene body in cells lacking INO80 and ISW2. These results suggest that RSC engages in a tug-of-war with INO80 and ISW2, which could lead to sharpening of the +1 position ^3^. H2A.Z on the other hand has no observable effect on +1 positioning, in line with prior results ^64^.

In the fourth process, remodelers of the ISWI, Chd1 and INO80 families space nucleosomes in gene bodies **(Fig. 8)**. They can readily override DNA-sequence encoded nucleosome spacing, particularly over genes that have extreme NRLs. The NRL distribution thereby considerably sharpens. In three quarters of all genes, however, nucleosomes end up again on positions with equally strong nucleosome affinities. DNA sequence therefore codetermines NRLs in most of the genome, and even ATP-dependent remodelers cannot level the thermodynamic landscape for nucleosomes.

**Fig. 8.**
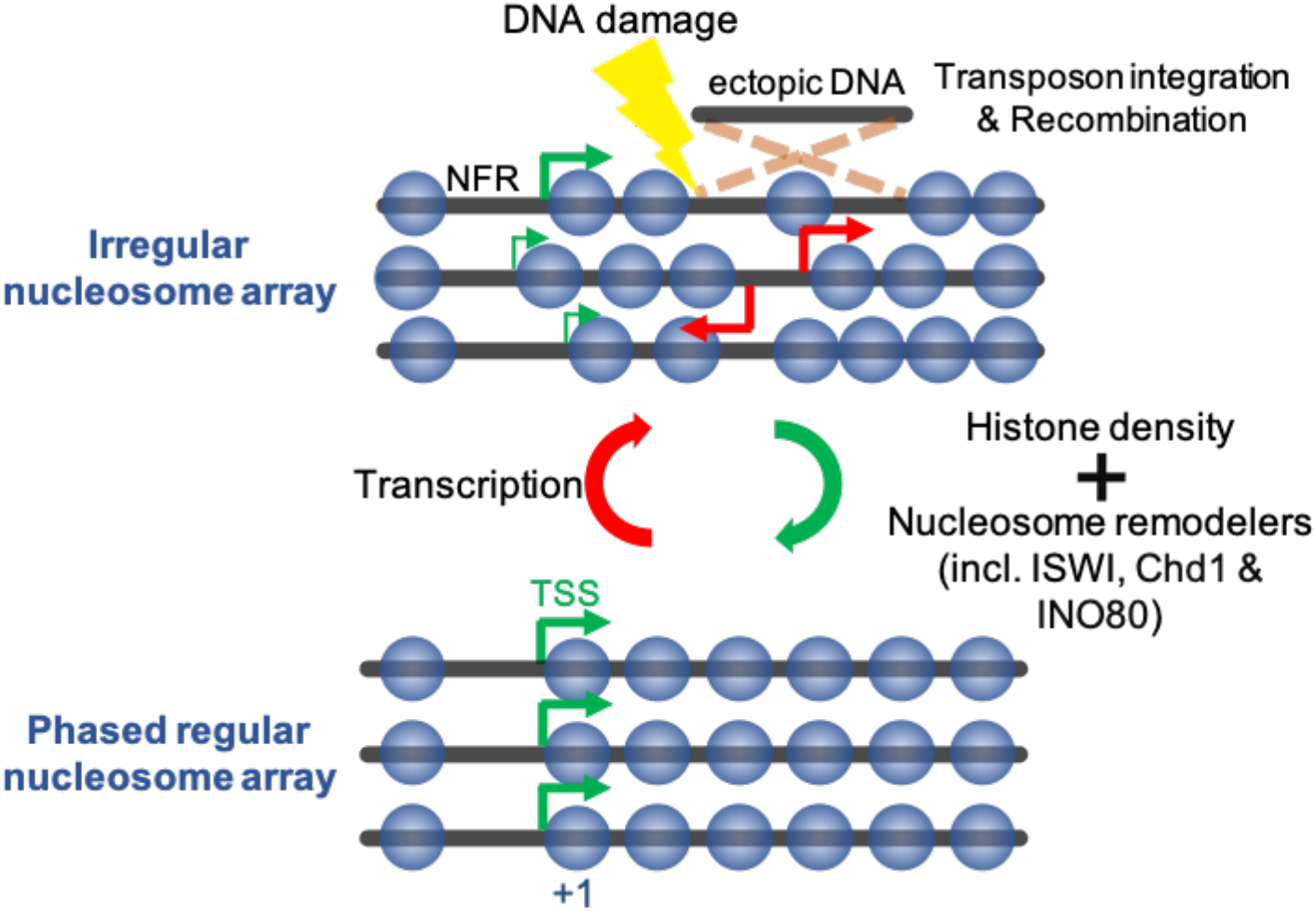
Model for regular nucleosome array organization. Transcription is strongly disruptive to nucleosome arrays and it takes ISWI-, Chd1- and/or INO80-remodelers to reinstate array architecture. Proper histone density is indispensable for spacing as neither DNA sequence-based nucleosome positioning nor the ‘clamping’ activity of remodelers suffices. Remodelers also position the +1 nucleosome, thereby sharpening the site of transcription initiation (green arrows). Regularly spaced nucleosome arrays prevent cryptic transcription (red arrows), DNA damage, ectopic recombination and transposon integration within the gene body.

Our nucleosome depletion experiments suggest that at least one ISWI or Chd1 remodeler possesses a clamping activity and can hold nucleosomes at a close distance even when nucleosome concentrations are diminished. Note that this result does not rule out the linker length equilibration mechanism ^14^. In fact, the clamping activity could be a manifestation of linker length equilibration provided that the remodeler can measure linker lengths only over short ranges. Nevertheless, the observed clamping activity is too weak to overcome the entropic forces acting on nucleosomes upon their depletion. The biogenesis of arrays therefore also requires WT-like nucleosome densities **(Fig. 1a)**.

We find that INO80 generates a broad NRL distribution peaking at 168 bp **(Supplementary Fig. 4c)**. This NRL falls short of the ∼200 bp linker length generated *in vitro* ^26^. The high cellular density of nucleosomes probably prevents INO80 from dispersing nucleosomes that far. When nucleosome densities naturally drop, however, for instance during DNA damage ^51^, INO80 may be free to induce a wider spacing.

The physiological NRL of ∼165 bp has been suggested to result from a competition between ISWI and Chd1 remodelers, with Chd1 promoting narrower and ISW1 wider spacing ^9^. Future models should also take INO80 into account. As each remodeler appears to prefer different, characteristic NRLs, we imagine that the cell could locally adjust nucleosome spacing by recruiting different spacing factors.

### The function of nucleosome arrays

Why do eukaryotic cells evenly space nucleosomes? Our findings indicate that even spacing between nucleosomes helps prevent cryptic transcription, probably by occluding cryptic TSSs, and thereby forcing the transcription machinery to generate proper, stable gene products **(Fig. 7h)**.

In addition, our data indicate that regularly spaced nucleosome arrays may have evolved to counteract integration of transposons **(Fig. 7e, f)**. Even spacing of nucleosomes could for example prevent occasional exposure of a large enough stretch of naked DNA that mobile elements or retroviruses would exploit to integrate into the genome. Similar considerations would explain our observation that TKO cells with their irregular nucleosome array structure suffer from higher homologous recombination rates than WT cells **(Fig. 7d)**.

Furthermore, we suggest that spacing remodelers have evolved to suppress genotoxicity from DNA damaging events. This model would explain why the DNA of cells lacking ISWI and Chd1 remodelers very quickly fragments in response to DNA-damaging agents **(Fig. 7b)**, and why genes with irregularly spaced nucleosomes are particularly prone to DNA double strand breaks **(Fig. 7c)**.

Protection against mobile elements, retroviruses, homologous recombination, cryptic transcription, and DNA damage may have been powerful evolutionary advantages that ensured retention of spacing remodelers throughout the eukaryotic domain.

## Methods

### Yeast strain generation

All yeast strains used in this study are derived from the W303 background and can be found in Supplementary Table 1. To validate gene deletions and tagging, all loci of interest were tested by PCR using oligonucleotides stated in Supplementary Table 2.

To generate a strain lacking *ISW1, ISW2* and *CHD1* in the anchor-away background, HHY170 ^30^ was mated to the strain YTT227 ^27^ and haploids were obtained via tetrad dissection. Other deletions in all anchor-away strains were done via direct transformation. For TBP and INO80 depletion experiments, the *SPT15* and *INO80* genes were C-terminally tagged with a FRB-GFP fusion construct. To obtain a TKO histone depletion strain, the strain DY5734 ^65^ was mated to YTT227 and haploids were obtained via tetrad dissection. The Ycp50 *HHT2-HHF2* plasmid was switched to pRS413 Gal1-10 *HHT2*-*HHF2* (pFMP519) via 5-FOA selection.

### Yeast growth conditions

To deplete RNA Pol II, INO80 or TBP, cells were grown OD 0.2-0.3 before addition of vehicle (90% ethanol, 10% Tween-20) or rapamycin (1 μg/ml final concentration, dissolved in vehicle).

For histone depletion experiments, cells were initially grown overnight to OD 1.0, washed with pre-warmed SC media without carbon source and dissolved in pre-warmed SC + 2% glucose media to OD 0.5. Cells were grown at 30°C for 3 h and harvested for MNase-seq.

### Yeast nuclei preparation and MNase digestion

Yeast nuclei were prepared largely as described ^66^. Strains were grown overnight to OD 0.8-1.0 in YPAD media. Cells were harvested (3000 x g, 8 min, 4°C) and washed once with cold water. The pellet was weighed (wet weight), resuspended in 2 volumes of preincubation solution (0.7 M ß-mercaptoethanol, 28 mM EDTA pH 8.0) and shaken at 30°C for 25-30 min. Cells were washed with 40 ml ice-cold 1 M sorbitol and resuspended in 5 volume of 1 M sorbitol, 5 mM ß-mercaptoethanol. 1 mg of freshly dissolved Zymolyase was added / g wet weight and incubated at 30°C for 20-30 min until the OD 600 reading was decreased to 80-90% of the initial OD. Cells were also checked under the microscope for appearance of 80-90% ghosts. Spheroplasts were collected by centrifugation (2500 x g, 5 min, 4°C) and washed with 40 ml ice-cold 1 M sorbitol. Spheroplast pellets were resuspended in 7 ml / g wet weight Ficoll buffer (18% Ficoll, 20 mM KH_2_PO_4_ pH 6.8, 1 mM MgCl_2_, 0.25 mM EGTA, 0.25 mM EDTA). Nuclei were centrifuged (12000 x g, 30 min, 4°C) and stored at −80°C.

For MNase digestion, nuclei were thawed on ice for 10 min, washed once with 8 ml MNase digestion buffer (15 mM Tris-Cl pH 7.5, 50 mM NaCl, 1.4 mM CaCl_2_, 0.2 Mm EGTA pH 8.0, 0.2 mM EDTA pH 8.0, 5 mM ß-mercaptoethanol), dissolved in 1 ml MNase digestion buffer and divided into 5 aliquots. An increasing amount of MNase (between 4-256 U / μl; dissolved in 10 mM HEPES-KOH pH 7.6, 50 mM NaCl, 1.5 mM MgCl_2_, 0.5 mM EDTA, 10% Glycerol, 1 mM DTT, 0.2 mM PMSF) was added, mixed and incubated for 20 min, 37°C. MNase digestion was stopped by adding 35 μl of quenching solution (10 mM EDTA, 1% SDS, 50 mM Tris-Cl pH 8.5). 300 μg Proteinase K was added and incubated at 37°C for 30 min. Afterwards, 70 μl of 5 M NaClO_4_ was added. The samples were extracted with Phenol: Chloroform: Isoamyl alcohol (25:24:1), then with Chloroform: Isoamyl alcohol (24:1) and precipitated with 100% ethanol. Pellets were washed with 70% ethanol and dissolved in TE buffer. RNA was removed by adding 10 μg RNase A and incubation at 37°C for 60 min. DNA was precipitated by adding 0.2 M NaCl (final concentration) and 0.7 volumes of Isopropanol, washed with 70% ethanol and dissolved in 50 μl TE buffer. To visualize the digestion degree, 25 μl of DNA was mixed with 3 μl of loading dye (0.5% Orange G, 50% glycerol) and electrophoresed on 1.7% low-melt agarose gel. Whenever possible, samples showing 70% mono-, 20% di-, 10% tri-nucleosome bands were used for high-throughput sequencing.

### Cell cycle profile

To measure cell cycle progression, 10^7^ cells were harvested, resuspended in 1 ml of 50 mM Tris pH 8.0, 70% ethanol and incubated overnight at 4°C. Cells were washed with 50 mM Tris pH 8.0 and incubated with 520 μl of RNase solution (500 μl of 50 mM Tris pH 8.0, 20 μl of 10 mg/ml RNase A) overnight at 37°C. Cells were treated with 220 μl Proteinase K solution (200 μl 50 mM Tris pH 8.0, 20 μl Proteinase K) for 30 min at 50°C and resuspended in 500 μl of 50 mM Tris pH 8.0 and sonicated for 15 sec (Bioruptor Pico). 15 μl of cells were stained with 285 μl of SYTOX solution (10 mM Tris pH 8.0, 1:10000 dilution, Cat# S7020 Life Technologies) and analyzed with BD FACSCanto or BD Fortessa (Core Facility Flow Cytometry Biomedical Center, LMU Munich).

### G1 cell cycle arrest

Cells were grown to OD 0.2 in YPAD media and alpha factor (10 μg/ml final concentration) added. After 1 h incubation, same amount of alpha factor was added, and cells were incubated for additional 1 hour. Cells were again supplemented with alpha factor (5 μg/ml final concentration), divided equally into two flasks and treated with either vehicle or rapamycin.

### RNA Pol II ChIP

Cells were grown to OD 0.4 in 500 ml YPAD media and treated with vehicle or rapamycin. Cells were crosslinked (1% formaldehyde, 10 min, 25°C) and quenched with 125 mM glycine. Cells were harvested, washed with ST buffer (20 mM Tris-HCl pH 7.5, 100 mM NaCl) + protease inhibitors (PI; 1 μg/ml Aprotinin, 1 μg/ml Leupeptin, 1 μg/ml Pepstatin A, 1 mM PMSF). Cell walls were disrupted in FA-lysis buffer (50 mM HEPES pH 7.5, 140 mM NaCl, 1 mM EDTA pH 8.0, 1% Triton X-100, 0.1% Na Deoxycholate, PI) by bead beating (Precellys 24 homogenenizer). Chromatin was fragmented to an average size of 500 bp using Diagenode Bioruptor Pico for 20-25 cycles of 30’’ on / 30’’ off in FA-lysis buffer + 0.25% SDS. SDS was diluted in the supernatant to 0.1% with FA-lysis buffer and incubated with 2 μg anti-Rpb3 antibody (Abcam 1Y26) for 2 h at 4°C. 100 μl of pre-washed Pan-Mouse IgG Dynabeads were added and incubated overnight at 4°C. Beads were washed twice with FA-lysis buffer, twice with FA-lysis + 360 mM NaCl buffer, twice with FA-wash 2 buffer (0.25 M LiCl, 0.5% NP40, 0.5% Na-deoxycholate, 1 mM EDTA, 10 mM Tris-Cl pH 8.0) and once with TE buffer. DNA was eluted in 50 μl TE buffer + 0.5% SDS at 65°C for 1 h. RNA was removed by with RNase A and DNA was de-crosslinked overnight by incubating with 10 μg Proteinase K at 65°C. Input DNA (1%) was treated the same in parallel. IP and input DNA were quantified by qPCR using Fast SYBR master mix (Life Technologies, Cat# 4385618). Primer sequences can be found in Supplementary Table 2.

### NGS library preparation and sequencing

We prepared sequencing libraries directly from the whole MNase digested samples, and not from mononucleosomal DNA, to avoid artifacts arising from imprecise gel extraction. DNA fragments longer than 500 bp were removed using AMPure size selection as follows: 300 ng DNA was diluted in 50 μl 0.1X TE buffer. 0.65X volumes of AMPure XP beads were added. 1.92X volumes of Ampure XP beads were added to the supernatant in a new eppendorf. Beads were washed twice with 500 μl freshly prepared 80% ethanol, and DNA was eluted in 30 μl 0.1X TE buffer. 50 ng of DNA was used for library preparation using NEBNext Ultra II DNA Library Prep kit for Illumina. Only 3 or 4 PCR cycles were performed to reduce potential bias arising from PCR for different fragment sizes in the library. Fragment lengths of ∼270-280 bp were consistently observed using the Bioanalyzer DNA kit. Libraries were sequenced on Illumina HiSeq for 50 cycles in paired-end mode.

### Western blot

Cells were grown overnight, reinoculated in fresh 10 ml YPAD to OD 0.1 and grown to OD 0.8. Extracts were prepared via the NaOH / TCA precipitation method. Blots were incubated overnight with anti-FLAG M2 (1:20000 dilution; Cat# F1804, Sigma-Aldrich) or H3 antibody (1:20000 dilution; Cat# ab1791, Abcam) or H4 antibody (1:2000 dilution; Cat# ab10158, Abcam) in 5% skimmed milk + PBS with 0.1% Tween-20. LI-COR IR secondary antibodies (1:10000 dilution; Cat# 926-68070, 926-32210, 926-32211) and Odyssey IR imaging system were used for visualization.

### Spotting assay

Cells were grown overnight to near saturation and OD 600 was measured in technical replicates by 1:10 dilution in water. Cells were diluted to OD 1.0 in 200 µl water and 5-fold dilutions were generated. 7 µl of these dilutions were spotted on YPAD supplemented with desired compound as necessary. Plates were incubated at 30°C for 3 days. Assay was performed twice using two independent colonies of the mutant strains.

### DNA damage assay

To test fragmentation of the genomic DNA, cells were grown overnight in 50 ml YPAD to OD 0.4 - 0.8. The log-phase cultures were diluted to OD 0.2 in 5 ml YPAD media and zeocin was added to 1 mg/ml final concentration. Cells were incubated at 30°C for 10 min with gentle shaking. Cells were harvested, washed with 5 ml ice-cold water and dissolved in 200 µl DNA extraction buffer (0.9 M Sorbitol, 50 mM Na-Pi pH 7.5, 140 mM ß-Mercaptoethanol) with 15 mM sodium azide. Cell walls were digested with Zymolyase (0.5 mg/ml, 15 min), followed by Proteinase K digestion (2 mg/ml, 15 min), both at 30°C. Phenol: Chloroform: Isoamyl alcohol (25:24:1) extraction was performed once, followed by ethanol precipitation. Pellets were dissolved in 100 µl TE buffer supplemented with 5 µg RNase A and incubated for 30 min at 30°C. Equal amounts of DNA were loaded on 0.6% (w/v) low-melt agarose gel in 1X TBE buffer and electrophoresed at 2 V/cm for 4 h. The gel was stained with ethidium bromide (0.5 µg/ml) in 1X TBE for 15 min. Samples were pipetted only three times with ends of pipette tips cut off to minimize fragmentation arising from shearing during pipetting.

### Ectopic recombination assay

The ectopic recombination rate was determined as described ^51^. An equal number of log phase cells (10^7^) were transformed. For insertions at the *ura3-1* locus, StuI digested pRS406 plasmid was used. For insertions at the *BAR1* locus, the HIS3 marker was amplified from pRS403 plasmid. As a negative control, undigested pRS406 or pRS403 plasmids (100 ng) were transformed. As a positive control, 100 ng of pRS416 or pRS413 plasmids were transformed in parallel. To calculate relative integration rates, the numbers of colonies obtained on StuI digested pRS406 or *HIS3* PCR product were first divided by the number of colonies obtained for their positive controls (pRS416 or pRS413, respectively). Normalized count for each replicate was divided by the normalized count observed in WT cell to obtain the relative integration rate.

### Cloning via Gibson assembly

To clone the *ARP8* gene including its native promoter and terminator, the *ARP8* locus was PCR amplified from genomic DNA. To delete the N-terminal region (amino acids 2-197) from *ARP8*, inverse PCR was performed on the plasmid with WT *ARP8*.

To clone the Gal1-10 *HHT2*-*HHF2* construct from the pRM102 ^28^ plasmid into the pRS413 vector, the Gal1-10 *HHT2*-*HHF2* construct was PCR amplified.

### ATAC-seq

A detailed ATAC-seq protocol for yeast ^67^ was generously provided by William Greenleaf and used with minor changes. Transposition reactions were performed in 25 µl tagmentation mix (1.25 µl Illumina Tagment DNA Enzyme (Cat# 20034197), 12.5 µl 2X Illumina Tagment DNA buffer, 11.25 µl water) for 15 min, 37°C with gentle shaking. DNA was purified and PCR amplified in 50 µl reaction (10 µl tagmented DNA, 10 µl water, 2.5 µl each Nextera index i5 and i7 primers and 25 μl NEBNext High-Fidelity 2X PCR Master Mix) initially for 5 cycles. Then, qPCR was performed on the amplified samples to calculate the minimum number of cycles (usually 4) required to avoid over-amplification. Libraries were size selected aiming for final fragment size 100 – 600 bp.

### Data analysis

All MNase-seq experiments were performed in biological replicates using two independent colonies, except for Rpb1 anchor-away and Nhp10 deletion in WT backgrounds, for which published datasets exist ^34,35,40^. Nuclei preparation and MNase digestion from two yeast colonies were performed on different days. Sequencing libraries were prepared in parallel for both colonies. Biological replicates were first analyzed separately and, in most cases, found highly consistent. For final analysis, bam files of biological replicates were merged for further downstream analyses.

### Demultiplexing, mapping and coverage

Fastq files from Illumina HiSeq were demultiplexed using Je demultiplex suite v1.0.6 with demultiplex-illu in paired-end mode. Sequences were mapped to *S. cerevisiae* sacCer3 R64-1-1 genome using Bowtie2 v2.2.9 with default settings, except -X 500, --no-discordant, --no-mixed options. Bam files were created using samtools v1.3.1 with minimum mapping quality 2 and mitochondrial (chrM) and rDNA reads (chrXII 451000:469000) were removed. Nucleosome dyad coverage and bigWig files were generated in R using rtracklayer v1.42.2, GenomicRanges v1.34.0 and GenomicAlignments v1.18.1 packages by taking the center of 140-160 bp fragments and resizing to 50 bp at the dyad center. All samples were sub-sampled to 10 million reads to generate an equal number of reads. The sequencing coverage was normalized to reads per million.

### Composite plot and heatmap

Genome coordinates and annotations for all genes, including the +1 nucleosome, were downloaded from ref ^68^. A matrix aligned to the +1 nucleosome was generated using coverageWindowsCenteredStranded function in tsTools v0.1.2 (https://github.com/musikutiv/tsTools). Composite plots were created from the aligned matrix by calculating the mean signal for each base pair of all genes and normalized to the mean signal of the desired window. Heatmaps were also generated using the aligned matrix.

### NRL and regularity score calculation

A MATLAB routine for calculating the NRL was generously provided by Răzvan Chereji and David Clark ^9^. We adapted it in R (https://github.com/musikutiv/tsTools) with some modifications. For each gene, the genomic region from a −200 to +800 bp window relative to the +1 nucleosome was used and smoothed with a 75 bp smoothing window. The smoothed nucleosome patterns were cross correlated with a theoretical periodic pattern of gaussian distributions with increasing repeat lengths. For each gene, the NRL was taken from the nucleosome pattern that resulted in the highest cross-correlation score. This cross-correlation score was used as an estimate for the array regularity with higher coefficients indicating higher array regularity.

### ATAC-seq analysis

Paired-end reads were processed and mapped to sacCer3 R64-1-1 genome using Bowtie2 v2.2.9 with default settings. A coverage vector was generated from the bam files taking fragments between 50 to 500 bp with equal number of reads (3 million) for each sample. A matrix aligned to the TSS and TTS ^68^ were generated from the coverage vector and gene bodies were re-scaled to 1000 bp.

For ATAC to array regularity correlation, reads mapping between TSS +100 bp and TTS −100 bp, and genes with mRNA longer than 300 bp and nucleosome occupancy between 0.78 and 0.88 ^61^ were considered for this analysis.

### Published datasets

RNA Pol II anchor-away, RSC and INO80 depletion and *nhp10Δ* MNase-seq datasets in the WT background were taken from ref ^3,34,35,40,47^, processed and plotted like datasets generated in this study. Rpb3 ChIP-seq datasets were taken from ref ^9^. Bam files with fragment lengths between 50 and 300 were used to calculate average RNA Pol II occupancy (IP / input) over each gene using the bamR package (https://github.com/rchereji/bamR).

### DANPOS +1 nucleosome calling and fuzziness analysis

Bam files with fragment lengths between 140 and 160 bp were used as input. Nucleosomes were called using dpos command with -jw 5, -q 200, -m 1 parameters in DANPOS v2.2.2 ^43^. The first nucleosome called after the NDR coordinate ^68^ of each gene was used as the +1 nucleosome and summit value for each gene was taken.

### Simulated statistical positioning after histone depletion

Simulations with 50000 DNA fragments between 2000 and 2500 bp in length were carried out in MATLAB (The Mathworks). The +1 nucleosome was positioned on one end of each DNA fragment using a gaussian function (sigma of 25 bp) that approximates MNase-seq derived values. All other nucleosomes were successively placed on a random DNA fragment at a random position, provided that this position was not occupied already by another nucleosome. Nucleosomes were modeled as hard spheres with a footprint of 146 bp. Random placement of nucleosomes continued until nucleosomes covered ∼51% of available DNA. After nucleosomes achieved this target density, 20% of all nucleosomes (picked at random, but excluding the +1 nucleosome) were dissociated again. The dissociated nucleosomes were placed on the DNA fragments again as above, and the dissociation and placement cycle was repeated another eight times. Three individual replicates of simulations were highly similar and simulation 1 is shown.

### Nucleosome affinity prediction using nuCpos

Nucleosome affinities were calculated with the Histone Binding Affinity (HBA) function in nuCpos ^48^, which provides a histone binding affinity score for a given 147 bp DNA sequence. The genome was divided into 147 bp sequences with a 1 bp sliding window step size. The HBA signal was aligned to +1 nucleosome ^68^ and smoothed using smooth.spline function (spar = 0.4). Smoothing using the rollmean (Zoo package) or sgolayfilt (signal package) functions yielded similar results. Only genes with discernible array structures were selected by filtering for array regularities >0.5 and NRLs between 150 and 200 bp. Each quartile represents the average signal of 786, 502 and 688 genes in WT, TKO and TKO - Pol II samples, respectively.

### Nucleosome positioning sequence (NPS) from Ioshikhes et al., 2006

NPS data for coordinates −931 to +528 relative to start codon was re-aligned to +1 nucleosome by calculating the difference between ATG codons and the +1 nucleosomes ^68^. Genes with array regularity >0.5 and NRLs between 150 and 200 bp were used. NPS data were smoothed using rollmean (Zoo package v1.8-5) with step size 51.

### Correlations of array regularity with cryptic TSSs, ds breaks and transposon integrations

To correlate cryptic TSS ^32^, transposase insertions ^53^ and Spo11- and topoisomerase 2-induced ds breaks ^69^ to array regularity, counts mapping in the region TSS+100 to TSS-100 bp were summed for each gene and divided by the analyzed length. Genes were sorted by array regularity and divided into quartiles. For transposase insertion data ^53^, only essential genes were included as insertions in essential genes were mostly lethal. Genes longer than 500 bp and nucleosome occupancy between 0.78 and 0.88 were used for these analyses.

## Acknowledgements

We thank Core Facility Flow Cytometry at Biomedical Center, LMU for FACS analysis; Philipp Korber and his lab for initial help and strains; Peter B. Becker for discussions; Alain Verreault and Jesper Q. Svejstrup for histone depletion strains; Răzvan Chereji and David J. Clark for sharing NRL calculation scripts; Slawomir Kubik and David Shore for sharing ChEC-seq datasets, B. Franklin Pugh for sharing NPS files; Frank C. P. Holstege for an Ino80-GFP-FRB construct; Boris Pfander for the *sgs1****Δ*** *exo1****Δ*** strain; Stefan Krebs and Helmut Blum (LAFUGA, Gene Center, LMU Munich) for sequencing. We are grateful to Silvia Härtel and Madhura Khare for technical help and the Müller-Planitz lab for discussions. A.K.S. acknowledges support from the DAAD for a predoctoral scholarship, and training by the IRTG SFB 1064 and CSHL Yeast Genetics and Genomics course. This work was supported by grants of the Deutsche Forschungsgemeinschaft (SFB1064 A07 and MU3613/3-1).

## Author contributions

Conceptualization: A.K.S., F.M.-P.;

formal analysis: A.K.S., T.Sc., T.St., F.M.-P.;

investigation: A.K.S., L.P.;

data curation: A.K.S., T.Sc., T.St.;

writing - original draft: A.K.S., F.M.-P.;

writing - reviewing and editing: A.K.S., F.M.-P.;

funding acquisition: F.M.-P.;

supervision: F.M.-P..

## Data availability

The next generation sequencing data has been deposited at the GEO under accession number GSE141007. All other relevant source data are provided in the manuscript.

## Code availability

All relevant code will be made available by the authors upon request.

## Biological materials

All materials generated in this study are available from the authors upon request.

## References

1. Lai, W.K.M. & Pugh, B.F. Understanding nucleosome dynamics and their links to gene expression and DNA replication. Nat Rev Mol Cell Biol 18, 548–562 (2017).

2. Jiang, C. & Pugh, B.F. Nucleosome positioning and gene regulation: advances through genomics. Nat Rev Genet 10, 161–72 (2009).

3. Kubik, S. et al.. Opposing chromatin remodelers control transcription initiation frequency and start site selection. Nat Struct Mol Biol 26, 744–754 (2019).

4. Weber, C.M., Ramachandran, S. & Henikoff, S. Nucleosomes are context-specific, H2A.Z-modulated barriers to RNA polymerase. Mol Cell 53, 819–30 (2014).

5. Groth, A., Rocha, W., Verreault, A. & Almouzni, G. Chromatin challenges during DNA replication and repair. Cell 128, 721–33 (2007).

6. Yadav, T. & Whitehouse, I. Replication-Coupled Nucleosome Assembly and Positioning by ATP-Dependent Chromatin-Remodeling Enzymes. Cell Rep 15, 715–723 (2016).

7. Vasseur, P. et al. Dynamics of Nucleosome Positioning Maturation following Genomic Replication. Cell Rep 16, 2651–2665 (2016).

8. Kubik, S. et al. Sequence-Directed Action of RSC Remodeler and General Regulatory Factors Modulates +1 Nucleosome Position to Facilitate Transcription. Mol Cell 71, 89–102 e5 (2018).

9. Ocampo, J., Chereji, R.V., Eriksson, P.R. & Clark, D.J. The ISW1 and CHD1 ATP-dependent chromatin remodelers compete to set nucleosome spacing in vivo. Nucleic Acids Res 44, 4625–35 (2016).

10. Gkikopoulos, T. et al. A role for Snf2-related nucleosome-spacing enzymes in genome-wide nucleosome organization. Science 333, 1758–60 (2011).

11. Ocampo, J., Chereji, R.V., Eriksson, P.R. & Clark, D.J. Contrasting roles of the RSC and ISW1/CHD1 chromatin remodelers in RNA polymerase II elongation and termination. Genome Res 29, 407–417 (2019).

12. Lieleg, C. et al. Nucleosome spacing generated by ISWI and CHD1 remodelers is constant regardless of nucleosome density. Mol Cell Biol 35, 1588–605 (2015).

13. Yamada, K. et al. Structure and mechanism of the chromatin remodelling factor ISW1a. Nature 472, 448–53 (2011).

14. Yang, J.G., Madrid, T.S., Sevastopoulos, E. & Narlikar, G.J. The chromatin-remodeling enzyme ACF is an ATP-dependent DNA length sensor that regulates nucleosome spacing. Nat Struct Mol Biol 13, 1078–83 (2006).

15. Stockdale, C., Flaus, A., Ferreira, H. & Owen-Hughes, T. Analysis of nucleosome repositioning by yeast ISWI and Chd1 chromatin remodeling complexes. J Biol Chem 281, 16279–88 (2006).

16. Udugama, M., Sabri, A. & Bartholomew, B. The INO80 ATP-dependent chromatin remodeling complex is a nucleosome spacing factor. Mol Cell Biol 31, 662–73 (2011).

17. Zhou, C.Y. et al. The Yeast INO80 Complex Operates as a Tunable DNA Length-Sensitive Switch to Regulate Nucleosome Sliding. Mol Cell 69, 677–688 e9 (2018).

18. Brahma, S., Ngubo, M., Paul, S., Udugama, M. & Bartholomew, B. The Arp8 and Arp4 module acts as a DNA sensor controlling INO80 chromatin remodeling. Nat Commun 9, 3309 (2018).

19. Knoll, K.R. et al. The nuclear actin-containing Arp8 module is a linker DNA sensor driving INO80 chromatin remodeling. Nat Struct Mol Biol 25, 823–832 (2018).

20. Kornberg, R.D. & Stryer, L. Statistical distributions of nucleosomes: nonrandom locations by a stochastic mechanism. Nucleic Acids Res 16, 6677–90 (1988).

21. Celona, B. et al. Substantial histone reduction modulates genomewide nucleosomal occupancy and global transcriptional output. PLoS Biol 9, e1001086 (2011).

22. Gossett, A.J. & Lieb, J.D. In vivo effects of histone H3 depletion on nucleosome occupancy and position in Saccharomyces cerevisiae. PLoS Genet 8, e1002771 (2012).

23. van Bakel, H. et al. A compendium of nucleosome and transcript profiles reveals determinants of chromatin architecture and transcription. PLoS Genet 9, e1003479 (2013).

24. Hughes, A.L., Jin, Y., Rando, O.J. & Struhl, K. A functional evolutionary approach to identify determinants of nucleosome positioning: a unifying model for establishing the genome-wide pattern. Mol Cell 48, 5–15 (2012).

25. Struhl, K. & Segal, E. Determinants of nucleosome positioning. Nat Struct Mol Biol 20, 267–73 (2013).

26. Krietenstein, N. et al. Genomic Nucleosome Organization Reconstituted with Pure Proteins. Cell 167, 709–721 e12 (2016).

27. Tsukiyama, T., Palmer, J., Landel, C.C., Shiloach, J. & Wu, C. Characterization of the imitation switch subfamily of ATP-dependent chromatin-remodeling factors in Saccharomyces cerevisiae. Genes Dev 13, 686–97 (1999).

28. Mann, R.K. & Grunstein, M. Histone H3 N-terminal mutations allow hyperactivation of the yeast GAL1 gene in vivo. EMBO J 11, 3297–306 (1992).

29. Kim, U.J., Han, M., Kayne, P. & Grunstein, M. Effects of histone H4 depletion on the cell cycle and transcription of Saccharomyces cerevisiae. EMBO J 7, 2211–9 (1988).

30. Haruki, H., Nishikawa, J. & Laemmli, U.K. The anchor-away technique: rapid, conditional establishment of yeast mutant phenotypes. Mol Cell 31, 925–32 (2008).

31. Shivaswamy, S. et al. Dynamic remodeling of individual nucleosomes across a eukaryotic genome in response to transcriptional perturbation. PLoS Biol 6, e65 (2008).

32. Lu, Z. & Lin, Z. Pervasive and dynamic transcription initiation in Saccharomyces cerevisiae. Genome Res 29, 1198–1210 (2019).

33. Zhang, Y. et al. Intrinsic histone-DNA interactions are not the major determinant of nucleosome positions in vivo. Nat Struct Mol Biol 16, 847–52 (2009).

34. Tramantano, M. et al. Constitutive turnover of histone H2A.Z at yeast promoters requires the preinitiation complex. Elife 5(2016).

35. Kubik, S. et al. Nucleosome Stability Distinguishes Two Different Promoter Types at All Protein-Coding Genes in Yeast. Mol Cell 60, 422–34 (2015).

36. Yen, K., Vinayachandran, V., Batta, K., Koerber, R.T. & Pugh, B.F. Genome-wide nucleosome specificity and directionality of chromatin remodelers. Cell 149, 1461–73 (2012).

37. Klein-Brill, A., Joseph-Strauss, D., Appleboim, A. & Friedman, N. Dynamics of Chromatin and Transcription during Transient Depletion of the RSC Chromatin Remodeling Complex. Cell Rep 26, 279–292 e5 (2019).

38. Papamichos-Chronakis, M. & Peterson, C.L. The Ino80 chromatin-remodeling enzyme regulates replisome function and stability. Nat Struct Mol Biol 15, 338–45 (2008).

39. Deniz, O., Flores, O., Aldea, M., Soler-Lopez, M. & Orozco, M. Nucleosome architecture throughout the cell cycle. Sci Rep 6, 19729 (2016).

40. Cutler, S., Lee, L.J. & Tsukiyama, T. Chromatin Remodeling Factors Isw2 and Ino80 Regulate Chromatin, Replication, and Copy Number of the Saccharomyces cerevisiae Ribosomal DNA Locus. Genetics 210, 1543–1556 (2018).

41. Tosi, A. et al. Structure and subunit topology of the INO80 chromatin remodeler and its nucleosome complex. Cell 154, 1207–19 (2013).

42. Watanabe, S. et al. Structural analyses of the chromatin remodelling enzymes INO80-C and SWR-C. Nat Commun 6, 7108 (2015).

43. Chen, K. et al. DANPOS: dynamic analysis of nucleosome position and occupancy by sequencing. Genome Res 23, 341–51 (2013).

44. Eustermann, S. et al. Structural basis for ATP-dependent chromatin remodelling by the INO80 complex. Nature 556, 386–390 (2018).

45. Brahma, S. et al. INO80 exchanges H2A.Z for H2A by translocating on DNA proximal to histone dimers. Nat Commun 8, 15616 (2017).

46. Ioshikhes, I.P., Albert, I., Zanton, S.J. & Pugh, B.F. Nucleosome positions predicted through comparative genomics. Nat Genet 38, 1210–5 (2006).

47. Ganguli, D., Chereji, R.V., Iben, J.R., Cole, H.A. & Clark, D.J. RSC-dependent constructive and destructive interference between opposing arrays of phased nucleosomes in yeast. Genome Res 24, 1637–49 (2014).

48. Kato, H., Shimizu, M. & Urano, T. Chemical map–based prediction of nucleosome positioning using the Bioconductor package nuCpos. bioRxiv (2019).

49. Alabert, C., Bianco, J.N. & Pasero, P. Differential regulation of homologous recombination at DNA breaks and replication forks by the Mrc1 branch of the S-phase checkpoint. EMBO J 28, 1131–41 (2009).

50. Morrison, A.J. Genome maintenance functions of the INO80 chromatin remodeller. Philos Trans R Soc Lond B Biol Sci 372(2017).

51. Hauer, M.H. et al. Histone degradation in response to DNA damage enhances chromatin dynamics and recombination rates. Nat Struct Mol Biol 24, 99–107 (2017).

52. van Attikum, H., Fritsch, O., Hohn, B. & Gasser, S.M. Recruitment of the INO80 complex by H2A phosphorylation links ATP-dependent chromatin remodeling with DNA double-strand break repair. Cell 119, 777–88 (2004).

53. Michel, A.H. et al. Functional mapping of yeast genomes by saturated transposition. Elife 6(2017).

54. Smolle, M. et al. Chromatin remodelers Isw1 and Chd1 maintain chromatin structure during transcription by preventing histone exchange. Nat Struct Mol Biol 19, 884–92 (2012).

55. Morrison, A.J. et al. INO80 and gamma-H2AX interaction links ATP-dependent chromatin remodeling to DNA damage repair. Cell 119, 767–75 (2004).

56. Baldi, S., Krebs, S., Blum, H. & Becker, P.B. Genome-wide measurement of local nucleosome array regularity and spacing by nanopore sequencing. Nat Struct Mol Biol 25, 894–901 (2018).

57. Cole, H.A., Ocampo, J., Iben, J.R., Chereji, R.V. & Clark, D.J. Heavy transcription of yeast genes correlates with differential loss of histone H2B relative to H4 and queued RNA polymerases. Nucleic Acids Res 42, 12512–22 (2014).

58. Schwabish, M.A. & Struhl, K. Evidence for eviction and rapid deposition of histones upon transcriptional elongation by RNA polymerase II. Mol Cell Biol 24, 10111–7 (2004).

59. Shen, C.H., Leblanc, B.P., Alfieri, J.A. & Clark, D.J. Remodeling of yeast CUP1 chromatin involves activator-dependent repositioning of nucleosomes over the entire gene and flanking sequences. Mol Cell Biol 21, 534–47 (2001).

60. Petesch, S.J. & Lis J.T. Rapid, transcription-independent loss of nucleosomes over a large chromatin domain at Hsp70 loci. Cell 134, 74–84 (2008).

61. Oberbeckmann, E. et al. Absolute nucleosome occupancy map for the Saccharomyces cerevisiae genome. Genome Res 29, 1996–2009 (2019).

62. Weiner, A., Hughes, A., Yassour, M., Rando, O.J. & Friedman, N. High-resolution nucleosome mapping reveals transcription-dependent promoter packaging. Genome Res 20, 90–100 (2010).

63. Kumar Singh, A. & Mueller-Planitz, F. Nucleosome positioning and spacing: from mechanism to function. J Mol Biol, 166847 (2021).

64. Hartley, P.D. & Madhani, H.D. Mechanisms that specify promoter nucleosome location and identity. Cell 137, 445–58 (2009).

65. Wittschieben, B.O., Fellows, J., Du, W., Stillman, D.J. & Svejstrup, J.Q. Overlapping roles for the histone acetyltransferase activities of SAGA and elongator in vivo. EMBO J 19, 3060–8 (2000).

66. Almer, A. & Horz, W. Nuclease hypersensitive regions with adjacent positioned nucleosomes mark the gene boundaries of the PHO5/PHO3 locus in yeast. EMBO J 5, 2681–7 (1986).

67. Schep, A.N. et al. Structured nucleosome fingerprints enable high-resolution mapping of chromatin architecture within regulatory regions. Genome Res 25, 1757–70 (2015).

68. Chereji, R.V., Ramachandran, S., Bryson, T.D. & Henikoff, S. Precise genome-wide mapping of single nucleosomes and linkers in vivo. Genome Biol 19, 19 (2018).

69. Gittens, W.H. et al. A nucleotide resolution map of Top2-linked DNA breaks in the yeast and human genome. Nat Commun 10, 4846 (2019).

