## Supplementary for "The biogenesis and function of nucleosome arrays"

SUPPLEMENTARY FIGURES

Supplementary Figure 1

**Histone depletion in WT and TKO cells.**

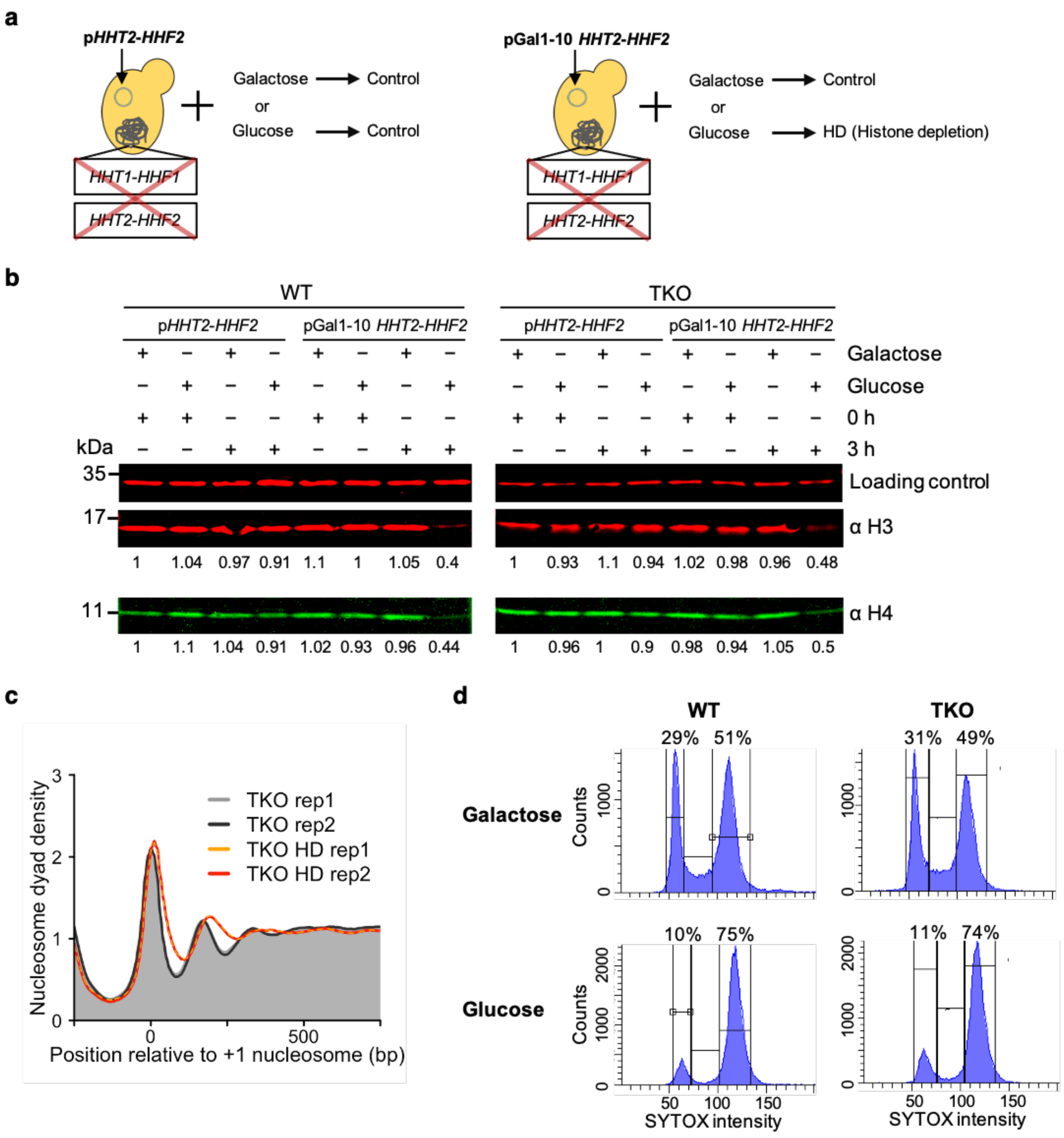


**a** Scheme of the histone depletion (HD) system [1]. Both genomic copies of histone H3 and H4 genes are deleted and replaced with a plasmid-borne copy of H3 and H4. On the plasmid, H3 and H4 either remain under the control of their native promoter (pHHT2-HHF2) or are put under control of the galactose-inducible Gal1-10 promoter (pGal1-10 HHT2-HHF2). Glucose-mediated repression of cells containing pGal1-10 HHT2-HHF2 leads to histone depletion. All other conditions serve as negative controls. **b** Glucose-mediated histone repression leads to reduction of H3 and H4 protein levels to ≥50% in WT and TKO cells. Numbers are relative expression levels between strains and conditions. These values represent the mean of two biological replicates. Values are normalized to a cross-reacting band seen with the anti-FLAG M2 antibody, which served as a loading control. Values were further normalized to H3 and H4 amounts measured in the left-most lane of each blot (pHHT2-HHF2 grown in galactose). **c** Composite plots showing nucleosome organization in TKO ad TKO HD cells in two biological replicates. **d** Cell cycle analysis upon histone depletion in WT and TKO cells. Flow cytometry profiles of cells grown in galactose or glucose containing minimal media for 3 h as in (**a**). Values represent percentage of cells in the G1 and G2/M phases.

**Supplementary Figure 2**

**RNA Pol II depletion increases array regularity in TKO cells.**

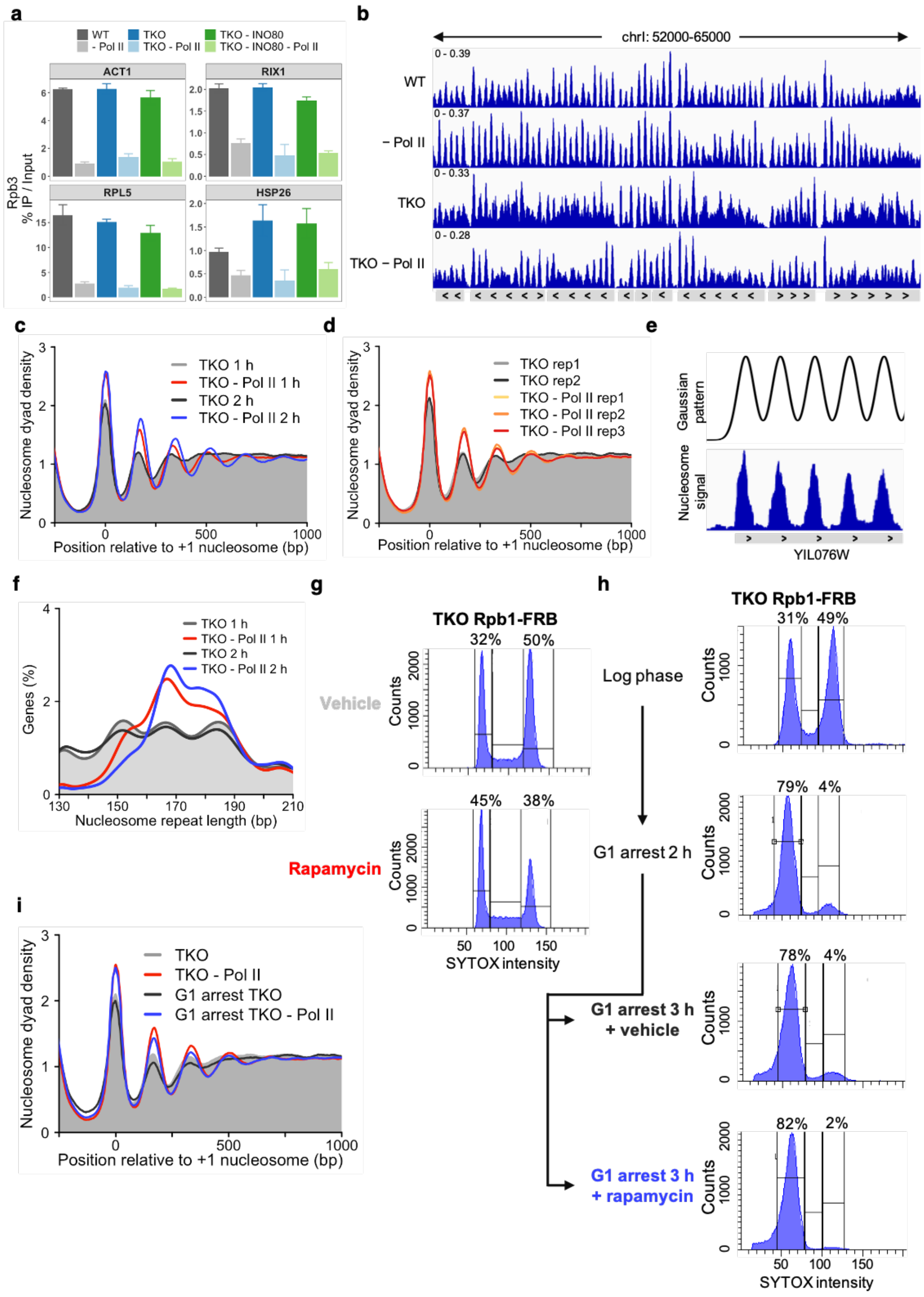

**a** Pol II is depleted from highly and lowly expressed genes upon addition of rapamycin in Rbp1-FRB tagged cells. ChIP-qPCR analysis of Rbp3 immunoprecipitation in WT, TKO and INO80 depleted TKO cells before and after Pol II depletion. Signals were normalized to input DNA used for Rbp3 immunoprecipitation of each strain. Errors: SD of three independent measurements. **b** Genome browser shot of MNase-seq in the indicated yeast strains. **c** Nucleosome organization upon Pol II depletion for 1 h and 2 h in TKO cells. TKO controls also harbored the FRB-tagged Rpb1 but were treated with vehicle. 1 h depletion data are replotted from **Fig. 2a**. **d** Biological replicates for **Fig. 2a**. **e** To obtain NRL and regularity over each gene, MNase-seq profiles over each gene are cross-correlated to Gaussian patterns of varying NRLs. The best-fitting Gaussian pattern provides an estimate of the NRL, and the correlation coefficient an estimate of the regularity. **f** NRL distribution of strains in c. Peak maxima after Pol II depletion are at 167 bp (1 h) and 168 bp (2 h), respectively. **g** Cell cycle analysis upon Pol II depletion in Rbp1-FRB tagged TKO cells. Flow cytometry profiles of cells treated with vehicle or rapamycin for 1 h. **h** Flow cytometry profiles of Rbp1-FRB tagged TKO cells before (log phase) and after treatment with alpha-factor for 2 h (G1 arrest 2h). Cells were then treated with vehicle or rapamycin for 1 h. Values represent percentages of cells in G1 and G2/M phases. **i** Nucleosome organization in cycling and G1 arrested Rbp1-FRB tagged TKO cells. Pol II was depleted for 1 h after a 2 h G1 arrest.

**Supplementary Figure 3**

**RNA Pol II depletion increases NRL and array regularity in WT cells.**

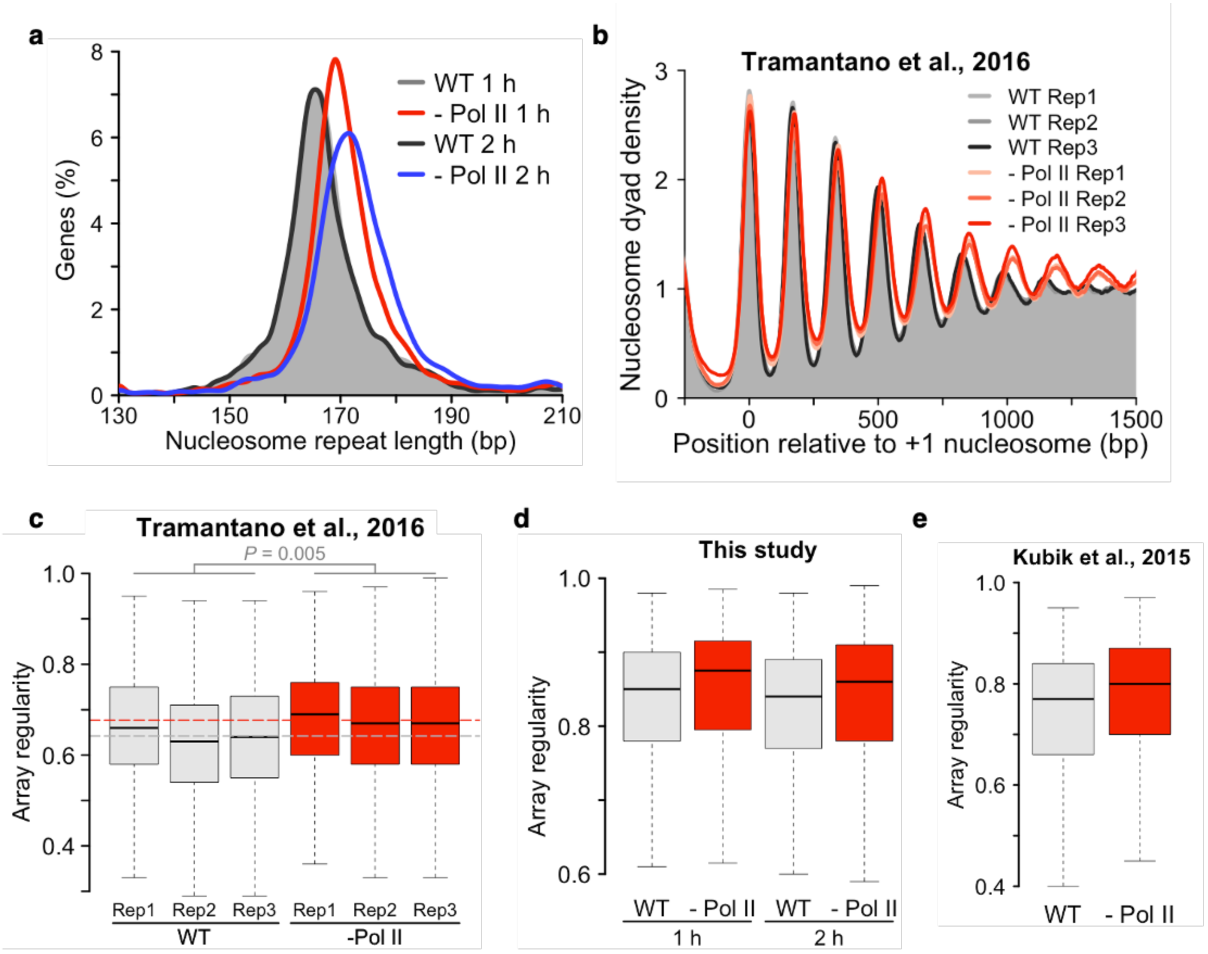


**a** NRL distribution before and after Pol II depletion in WT cells. Peak maxima are at 169 bp (1 h depletion) and 172 bp (2 h), respectively. Control samples are Rpb1-FRB tagged and vehicle-treated for 1 h or 2 h with peak maxima at 166 bp and 165 bp, respectively. **b** Gene averaged nucleosome organization in Pol II depleted cells. Data is from [2]. **c** Median array regularity increases upon Pol II depletion in WT cells. Boxplots represent array regularity in individual replicates (Rep) from [2]. *P* value represent statistical analyses performed with two-tailed paired Welch's t-test on the mean values of three replicates. **d, e** Same as (**c**) but for single replicates from this and a prior study [3]. Lack of replicates precluded calculation of *P* values.

**Supplementary Figure 4**

**The INO80 complex helps generate phased regular nucleosome arrays.**

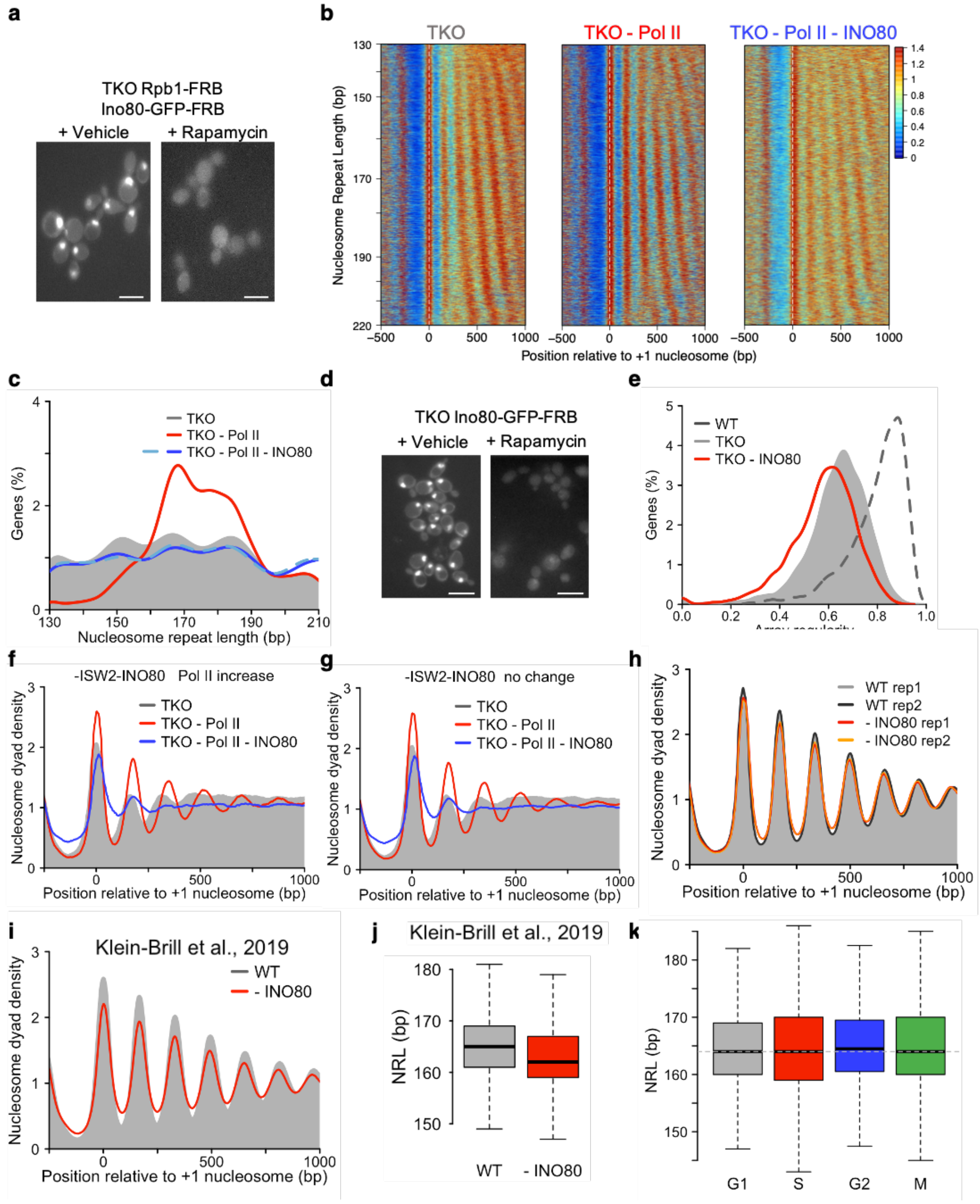


**a** Live-cell imaging of GFP-FRB-tagged INO80 shows its depletion from the nucleus. Treatment time 2 h. Scale bar 5 μm. **b** Heatmaps showing nucleosome organization upon Pol II- or combined Pol II- and INO80-depletion for 2 h in TKO cells. The TKO control is a 2 h vehicle-treated Rpb1-FRB TKO strain. Genes are sorted by NRL measured in the TKO control. **c** NRL distribution of samples in (**b**). **d** Live cell imaging for TKO Ino80-GFP-FRB cells. Treatment time 1.5 h. **e,** Array regularity distribution in TKO cells depleted for INO80. **f** Average nucleosome organization over 1592 genes that experience increased Pol II occupancy upon ISW2 and INO80 depletion, as identified in [4]. **g** Same as (**f**), but for genes that do not show increased Pol II levels (3330). **h, i** Nucleosome organization upon INO80 depletion in WT cells (**h**, 1.5 h anchor-away; **i**, 1 h auxin-induced degradation from [5]). **j,** NRL distribution upon INO80 depletion in WT cells. Data from [5]. **k,** NRL distribution in cells during G1, S, G2 and M cell cycle phases. Data from [6].

**Supplementary Figure 5**

**Role of INO80 domains and subunits in nucleosome organization.**

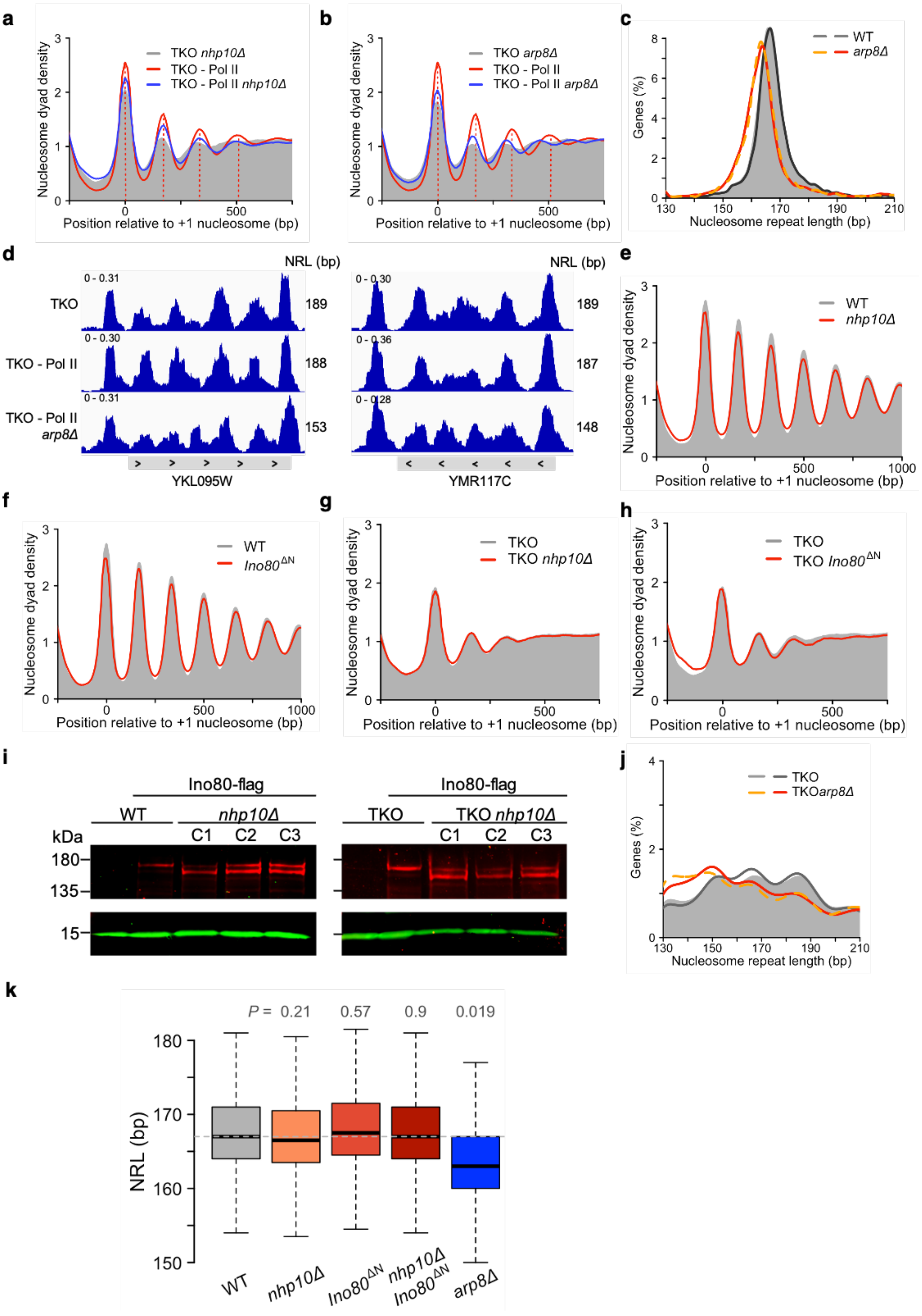

**a** Deletion of *NHP10* does not alter the position of nucleosome peaks that arise upon Pol II depletion (1 h) in the indicated strains (dashed lines). **b** Same as **a**, but for *arp8Δ*. The NRL visibly shortened upon *arp8Δ* deletion. **c** NRL distribution in WT and *arp8Δ* cells. Peak maxima are at 166 bp and 163 bp, respectively. **d** IGV browser shots of the indicated strains. *ARP8* deletion shortened the nucleosome-to-nucleosome distance. **e** Deletion of *NHP10* does not alter the NRL and nucleosome arrays. Reanalysis of published data also show negligible effects (not shown) [7]. **f** Deletion of 300 N-terminal amino acids of Ino80 does not affect the genome-wide nucleosome organization. **g, h** The nucleosome organization remains similar upon deletion of *NHP10* or Ino80's N-terminus in TKO cells. **i** Western blots showing degradation of Ino80 ATPase upon *NHP10* deletion in WT and TKO backgrounds. The Ino80 ATPase was C-terminally FLAG-tagged where indicated. Three independent *nhp10Δ* clones (C1, C2 and C3) were tested. **j** Deletion of *ARP8* shifts the NRL distribution to lower values in TKO cells. **k,** Box plots of measured NRLs in the indicated single and double mutants in otherwise WT cells. Horizontal line is at 167 bp. *P* values are from a two-tailed paired Welch's t-test on the mean values of two replicates relative to WT.

**Supplementary Figure 6**

**DNA sequence influences the NRL over genes.**

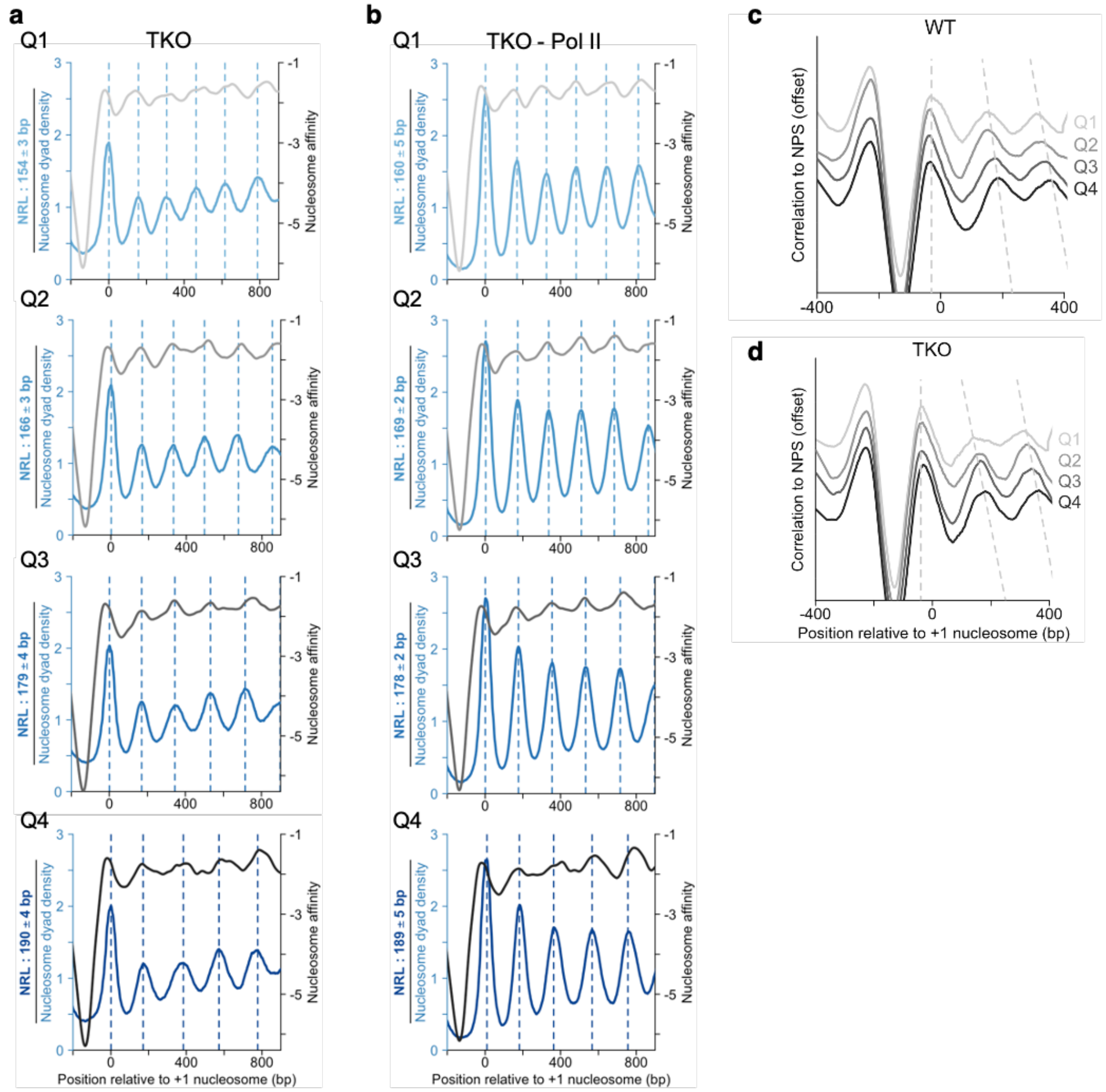


**a** Comparison of MNase-seq data from TKO cells with predicted nucleosome affinities. Genes were divided into quartiles according to their NRL. Mean values and SD of NRLs are provided on the left of each quartile. **b** Same as (**a**), but for Pol II-depleted TKO cells. **c, d** An alternative nucleosome prediction algorithm[8] (Nucleosome Positioning Sequence or NPS) reproduces results obtained in **Fig. 6d, e** respectively.

**Supplementary Figure 7**

**Regular nucleosome affect the function of the genome.**

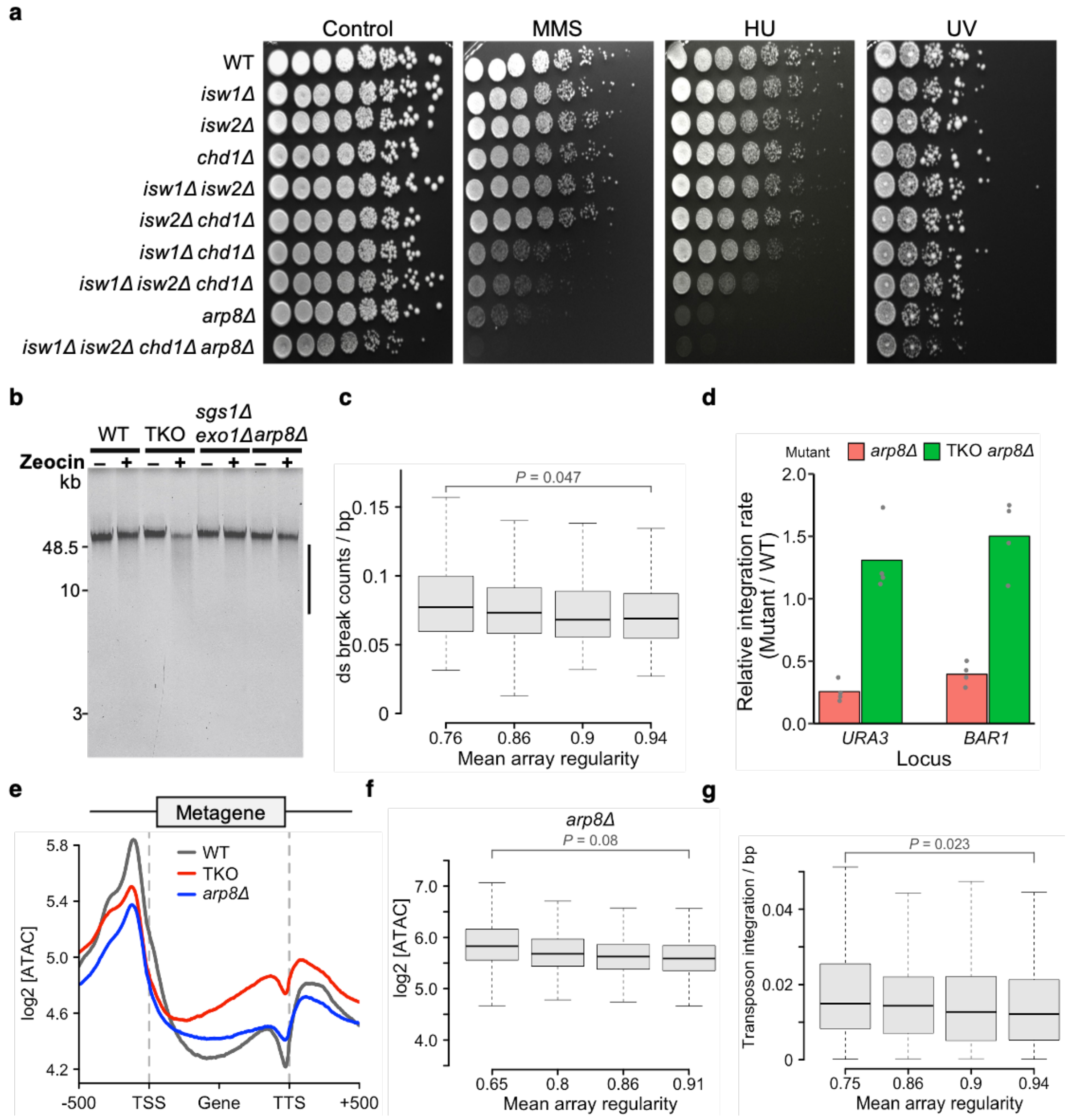


**a** Growth assay for indicated yeast strains on YPAD with or without Methyl methanesulfonate (MMS; 0.05%), or Hydroxyurea (HU; 200 mM). UV–treatment was 200 J/m$^2$. **b** Replicate from **Fig. 7b**. **c** Topoisomerase 2-induced DNA ds breaks that naturally occur during meiosis [9] enrich in genes with low array regularity. **d** Homologous recombination tested at two genomic loci (*URA3*, *BAR1*) in indicated mutants. Dots indicate four individual replicates. **e** ATAC-seq replicate (see **Fig. 7e**). **f** Array regularity does not significantly correlate with ATAC-seq signal in the gene bodies of *arp8Δ* cells. **g,** Ectopically induced transpositions *in vivo* anti-correlate with array regularity. Transposition data is from [10]. *P* values in (**c**, **f, g**) represent statistical analyses performed with two-tailed paired Welch's t-test on the mean values of two replicates.


1. Mann, R.K. & Grunstein, M. Histone H3 N-terminal mutations allow hyperactivation of the yeast GAL1 gene in vivo. *EMBO J* **11**, 3297-306 (1992).
2. Tramantano, M. et al. Constitutive turnover of histone H2A.Z at yeast promoters requires the preinitiation complex. *Elife* **5**(2016).
3. Kubik, S. et al. Nucleosome Stability Distinguishes Two Different Promoter Types at All Protein-Coding Genes in Yeast. *Mol Cell* **60**, 422-34 (2015).
4. Kubik, S. et al. Opposing chromatin remodelers control transcription initiation frequency and start site selection. *Nat Struct Mol Biol* **26**, 744-754 (2019).
5. Klein-Brill, A., Joseph-Strauss, D., Appleboim, A. & Friedman, N. Dynamics of Chromatin and Transcription during Transient Depletion of the RSC Chromatin Remodeling Complex. *Cell Rep* **26**, 279-292 e5 (2019).
6. Deniz, O., Flores, O., Aldea, M., Soler-Lopez, M. & Orozco, M. Nucleosome architecture throughout the cell cycle. *Sci Rep* **6**, 19729 (2016).
7. Cutler, S., Lee, L.J. & Tsukiyama, T. Chromatin Remodeling Factors Isw2 and Ino80 Regulate Chromatin, Replication, and Copy Number of the Saccharomyces cerevisiae Ribosomal DNA Locus. *Genetics* **210**, 1543-1556 (2018).
8. Ioshikhes, I.P., Albert, I., Zanton, S.J. & Pugh, B.F. Nucleosome positions predicted through comparative genomics. *Nat Genet* **38**, 1210-5 (2006).
9. Gittens, W.H. et al. A nucleotide resolution map of Top2-linked DNA breaks in the yeast and human genome. *Nat Commun* **10**, 4846 (2019).
10. Michel, A.H. et al. Functional mapping of yeast genomes by saturated transposition. *Elife* **6**(2017).
